# Chromosome topology gates productive RecA homology search

**DOI:** 10.64898/2026.07.20.739429

**Authors:** Miloš Tišma, Sourav Bhattacharyya, Seungwoo Chang, Taekjip Ha, Anjana Badrinarayanan, Joseph J. Loparo

**Affiliations:** Department of Biological Chemistry and Molecular Pharmacology, Blavatnik Institute, Harvard Medical School, Boston, MA 02115, USA; Howard Hughes Medical Institute and Program in Cellular and Molecular Medicine, Boston Children’s Hospital, Boston, MA 02115, USA; National Centre for Biological Sciences, Tata Institute of Fundamental Research, Bengaluru 560065, India

## Abstract

In homologous recombination, DNA repair depends on recombinase filaments finding homologous templates on chromosomes whose topology is continually remodeled by replication and transcription. How dynamic chromosome topology affects homology search and DNA repair remains unknown. By combining live-cell imaging of RecA-mediated homology search with single-molecule assays on topologically-defined DNA substrates, we show that DNA supercoiling acts as a selectivity filter for productive homology search. In *Caulobacter crescentus* cells, supercoiling is dispensable for the filament movement required for homology search, but required for homology target capture and repair. *In vitro*, filaments transiently sample both relaxed and supercoiled DNA, yet selectively commit to capture only on negatively supercoiled targets. Mechanistically, we find an extreme kinetic preference (~100-fold) of filaments towards negatively supercoiled DNA targets; this topological preference is independent of DNA compaction, pins plectonemes at the capture site, and is conserved across diverse bacterial RecA homologs. Our findings establish that chromosome topology physically regulates homology search and DNA repair.

## Introduction

Homologous recombination (HR) preserves genome integrity by locating an intact homologous sequence and using it as a template to repair broken DNA (*1*). During DNA repair, HR must solve two coupled problems: searching for a short homologous sequence on the chromosome and converting that encounter into a stable repair intermediate. Throughout this search, the chromosome is not a passive substrate but a dynamically supercoiled polymer continuously reshaped by replication, transcription, protein binding, and topoisomerases (*2*).

Central to HR are RecA/RAD51-family recombinases, conserved ATPases that catalyze homology search and strand exchange (*3*). Upon DNA damage, DNA processing by the HR machinery generates a 3′ single-stranded DNA (ssDNA) overhang (*4–7*). In bacteria, RecA assembles on this overhang to form a nucleoprotein filament termed the “presynaptic filament” (*8, 9*). This filament then searches the chromosome (*10*) powered by SMC-like proteins (*11*), and RecA’s own ATPase activity (*12, 13*), to ensure homologous target capture and downstream processes that result in high-fidelity repair. While searching for homology, the RecA filament must sample a vast pool of non-homologous sequences to identify and bind its complementary match.

Models for efficient homology search span multiple length scales, from structural insights involving “base flipping” (*14*) to cellular-scale models of genome-wide homology sampling by RecA filaments (*10*). Two models have been particularly influential in this regard. The ‘intersegmental transfer’ model (*15*) posits that DNA compaction enables rapid sampling of distant segments by a long RecA filament, while the ‘reduced-dimensionality search’ model (*10*) proposes that extended RecA filaments, positioned across the bacterial cell, can sample many chromosomal sites simultaneously. Additional factors including ssDNA binding (SSB) proteins (*16, 17*), DNA polymerases (*18*) and DNA topology in plasmids (*19–21*) have been implicated in facilitating homology search. Despite this, the factors that determine how transient sampling of the chromosome becomes productive target pairing competent for repair remains unresolved.

Single-molecule approaches have been especially powerful for studying homologous recombination because they resolve transient intermediates, mechanistic steps, and heterogeneous outcomes that are obscured in ensemble measurements (*8, 15, 22–24*). However, given the challenges of introducing defined supercoiling levels in DNA substrates, previous single-molecule studies of homology search have been largely restricted to relaxed substrates. Similarly, direct visualization of homology search in living cells has been unavailable in most systems and has only recently been achieved using specialized live-cell imaging designs (*11*). As a result, it remained unclear whether DNA topology is merely a structural consequence of chromosome organization or an active regulator of homology search and DNA repair.

Here we address how DNA topology of the chromosome impacts homology search. By developing a single-molecule assay in which target DNA supercoiling can be precisely tuned to physiological levels and combining it with live-cell imaging of homology search in *Caulobacter crescentus*, we show that negative supercoiling is the primary physical feature enabling productive homology search. Both *in vitro* and *in vivo*, lack of DNA supercoiling renders the homology search and capture unsuccessful. Thus, our results establish DNA supercoiling as a topological selectivity filter that links intrinsic chromosome organization to faithful DNA repair.

## Results

### Single-molecule assay for homology search on topologically defined DNA

To precisely study the effect of DNA topology on homology search, we developed a topology-controlled single-molecule assay that directly tests how supercoiling affects successful RecA homology search. Specifically, we combined a surface-based single-molecule assay (*25, 26*) with an analytical model which allowed us to precisely tune supercoiling density of target DNA molecules, while simultaneously monitoring both DNA and RecA-ssDNA dynamics.

We generated supercoiled DNA substrates by attaching a 42kb DNA molecule to the glass surface using multiple biotin-streptavidin attachment points close to the DNA ends (*25, 26*) (Fig. 1A, see Methods for details). Multiple surface attachments prevent free rotation around the central axis of DNA and the relaxation of torsional strain. We introduced defined levels of DNA supercoiling (Fig. S1A-B) by modulating the concentration of DNA intercalating dye SYTOX Orange (SxO). As SxO induces changes in base pair distance and twist angle (*25, 27*), its binding results in underwound DNA molecules which are torsionally constrained on the surface (Fig. S1A). Subsequent dye washout in low-SxO buffer traps this underwound state as negative writhe, producing negatively supercoiled plectonemes that cannot relax by free rotation (Fig. S1A, B). Roughly 20% of surface-attached DNAs were torsionally constrained (Fig. S1C) with typical bright foci resulting from plectonemes diffusing along the DNA length (*25*) (Fig. S1D). This internal mixture of constrained and relaxed molecules on the surface allowed us to directly compare homology search dynamics between supercoiled and relaxed molecules within the same experiment.

**Figure 1.**
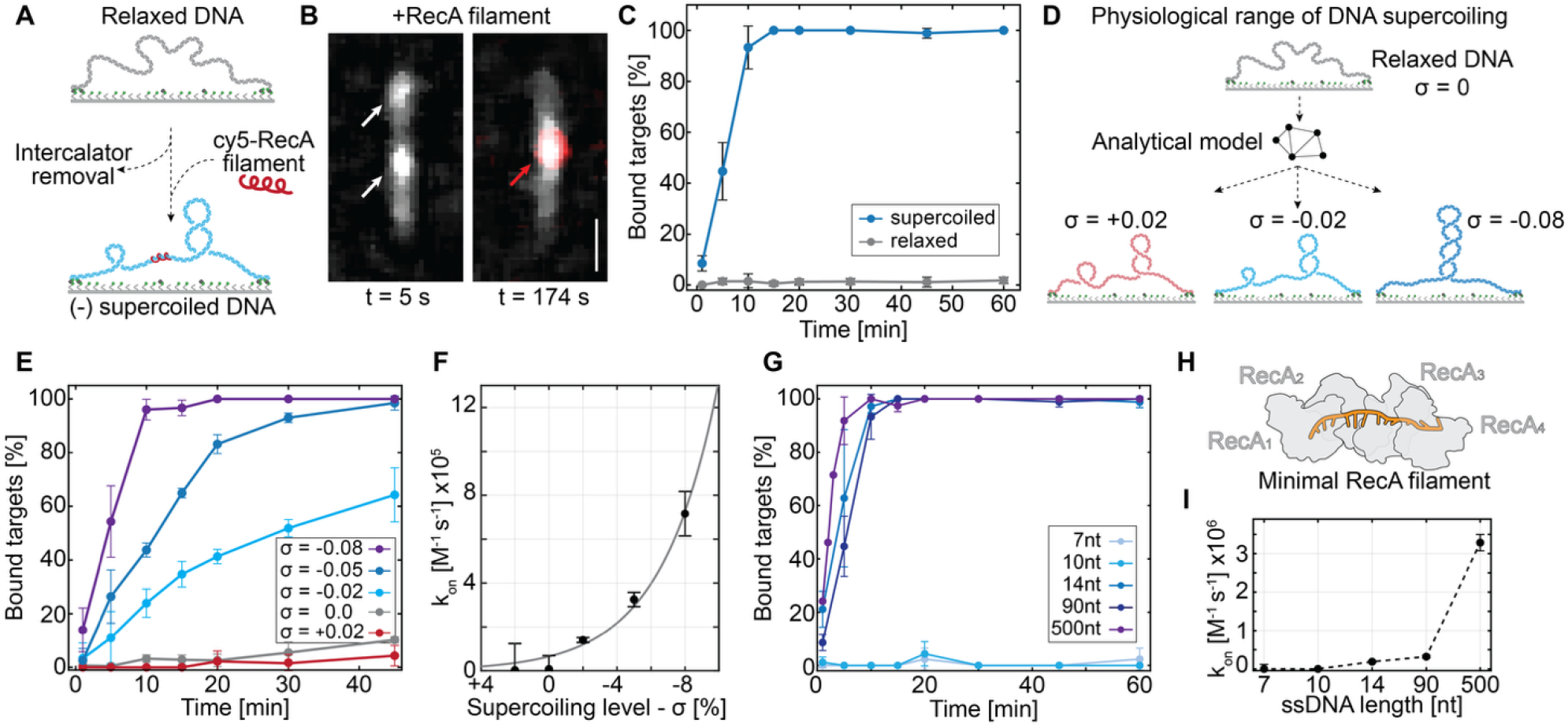
RecA-ssDNA presynaptic filaments strongly prefer negatively supercoiled DNA targets. **A)** Schematic representation of single-molecule surface assay. Topologically constrained dsDNA molecules have induced supercoiling while being tethered to the glass surface. Pre-assembled cy5-labelled RecA filaments are added to the flow cell to follow homology search on surface bound dsDNA target. **B)** Fluorescence images of dsDNA targets before (left, t = 5s) and after (right, t = 174 s) binding of Cy5-labelled presynaptic filament. White arrows point to observed plectoneme structures on supercoiled dsDNA. Red arrow points to bound Cy5-ssDNA-RecA presynaptic filament. Scale bar = 2 µm. **C)** Timecourse experiments tracking ssDNA-RecA filament (8 nM) binding to surface tethered DNAs. Supercoiled (blue, n = 642) and nicked (gray, n = 1256) molecules were directly compared within the same flow cell. Error bars represent standard deviation for each time point measurement (n=3). **D)** Introducing precise physiological supercoiling density (σ) into surface-bound DNA molecules by manipulating SxO intercalator concentrations, and an analytical model to calculate the resulting supercoiling density. **E)** Timecourse experiments on supercoiled molecules in the presence of 8 nM ssDNA-RecA (n_(_σ=+0.02) = 298, n_(_σ=0) = 1281, n_(_σ=−0.02) = 319, n_(_σ=−0.05) = 382, n_(_σ=−0.08) = 384). Error bars represent standard deviation for each time point measurement. **F)** Binding rates (k_on_) calculated by fitting the curves shown in E) with a pseudo-first order kinetic binding model. Error bars represent the standard error of the fitted pseudo-first-order rate constant. Exponential model fitted to the experimentally determined k_on_ values represented in gray line (k_on(_σ_=0)_ = 6.54 × 10^4^ M^−1^s^−1^, ß = 2.54, R^2^ = 0.983). **G)** Timecourse experiments using different invasion strand (ssDNA) lengths. (n_(7nt)_ = 330, n_(10nt)_ = 505, n_(14nt)_ = 532, n_(90nt)_ = 642 (same data as Fig. 1D), n_(500nt)_ = 603). Error bars represent standard deviation for each time point measurement. **H)** Cartoon depiction of the “minimal RecA filament” comprised of 4xRecA monomers (gray) and 14 nt ssDNA (orange). **I)** Binding rates (k_on_) calculated by fitting the curves shown in G) with a pseudo-first order kinetic binding model. Error bars represent the standard error of the fitted pseudo-first-order rate constant. Kinetic rates: k_on_ (14 nt) = 1.87 × 10^5^ ± 2.3 × 10^4^, k_on_ (90 nt) = 3.18 × 10^5^ ± 3.32 × 10^4^, k_on_ (500 nt) = 32.9 × 10^5^ ± 21.4 × 10^4^.

To observe homology search on our supercoiled substrates, we assembled *Escherichia coli* RecA filaments on a 90 nt Cy5-labelled ssDNA “invasion strand” in the presence of ATPγS. The invasion strand was fully complementary to a sequence positioned in the middle of the surface-tethered 42 kb target DNA molecule. We then slowly flowed these filaments over the surface-bound DNA targets and observed real-time homology search and homology pairing (Fig. 1B). Our assay allowed us to simultaneously observe presynaptic filaments (Cy5-ssDNA-RecA) and DNA plectonemes that form on supercoiled DNA (Fig. 1B).

### DNA topology determines efficient homology binding

Upon addition to the flow cell, RecA-ssDNA filaments bound rapidly and efficiently to negatively supercoiled DNA molecules (Fig. 1C). Such RecA filament binding was long-lived, stable and localized to the homologous region (see later). Within 15 min, 100% of supercoiled DNAs (85/85), were bound by RecA-ssDNA filaments (Fig.1C). This was in stark contrast to relaxed DNA molecules which showed essentially no binding in the same time period (5/543). Even after 1h, less than 3% of relaxed DNA molecules (14/1256) had bound filaments (Fig. 1C). Stable binding remained dependent on ATPγS and elevated Mg^2+^ concentration, consistent with previous work on plasmids(*28, 29*), and was strongly impaired at Mg^2+^ concentrations below 5 mM (Fig. S1E-G). By contrast, SSB protein did not considerably affect RecA filament binding kinetics (Fig. S1H).

Next, we precisely tuned the level of supercoiling to physiological ranges reported for *E. coli* (supercoiling density - σ ~ −0.06, or −6%)(*30*) and measured the effect on homology binding kinetics (Fig. 1D). To do this we developed an analytical model (*27*) that allowed us to determine the SxO concentration needed during surface tethering to generate average supercoiling densities consistent with cellular levels (+0.02 ≥ σ ≥ −0.08 (*30*), Fig. S2A-C). Notably, highly negatively supercoiled molecules appeared visibly stiffer and closer to the glass surface, consistent with increased tension (Movie S1). Positively supercoiled and relaxed DNA molecules showed negligible RecA filament binding, while even modest negative supercoiling (σ = −0.02) supported efficient filament binding (Fig. 1E). Further increases in negative supercoiling (σ = −0.05 and −0.08), progressively increased RecA filament binding rates (Fig. 1E). We quantified the binding rates at all supercoiling levels and observed the highest k_on_ rate for supercoiling density of σ = −0.08 at 7.1 ± 1.1 × 10^5^ M^−1^s^−1^ (Fig. 1F, S2D). An exponential model reproduced the experimental k_on_ rates with excellent agreement in the negatively supercoiled regime (Fig. 1F - gray line), however, the zero-torque rate (k_on(_σ =0)) predicted ~65% of relaxed DNA molecules would show binding during our experiments. This is a dramatic overestimate of the <10% of relaxed molecules bound by RecA filaments that we observed (Fig. 1E). Such discrepancy suggests that stable binding on relaxed DNA is suppressed, likely by energy required for duplex opening, or complex stability after target binding - at the commitment step. Overall, our data show that negative supercoiling is essential for a successful homology search and stable homology binding.

Successful homology search during HR requires presynaptic filaments to discriminate homologous from non-homologous sequences with high fidelity, a process shown to depend on homology length and low mismatch tolerance(*31–33*). As negative supercoiling is essential for stable RecA filament binding, we next asked whether our system retained dependence on filament length and sequence homology.

We first varied lengths of fully homologous filaments from 500 nt, 90 nt, down to 14 nt, 10 nt, and 7 nt. The latter three were selected to probe the minimal filament capable of stable homology search; given the structural consensus of ~3 nt/monomer(*34*) these correspond to tetrameric, trimeric, and dimeric RecA filaments, respectively. We observed efficient, stable binding at 14 nt invasion strand, corresponding to tetrameric RecA filaments, whereas 10 nt and 7 nt filaments showed no detectable binding events (Fig. 1G). While prior reports suggested that RecA filaments could stably form as dimers on a 6 nt ssDNA(*35*), under our supercoiling-calibrated conditions a stable minimal filament consisted of four RecA monomers on 14 nt ssDNA (Fig. 1H). Longer RecA filaments exhibited progressively higher binding rates, with 90 nt and 500 nt ssDNA filaments binding with ~1.7-fold and ~17.5-fold higher k_on_-rates, relative to 14 nt minimal RecA filament (k_on_ = 1.87 ± 0.23 × 10^5^ M^−1^s^−1^) (Fig. 1I).

We next asked whether homology search on supercoiled DNA substrates retained sequence dependence during homology recognition. For both 14 nt and 90 nt filaments, incomplete homology reduced binding relative to fully homologous RecA filaments (Fig. S3). For 90 nt filaments, randomized sequences of similar GC content associated with 2- to 8-fold slower kinetics and incomplete saturation during our experimental window (Fig. S3A-C), likely due to 8-9 nt microhomology across the 42 kb target(*24, 36*). In contrast, C-rich or T-rich 14 nt filaments lacking any microhomology stretches longer than 6 nt abolished binding entirely, while fully and partially homologous (8 out of 14 nt) short filaments showed comparable kinetics despite differences in homology length (Fig. S3D-G). Overall, RecA filaments retained high binding specificity in our assay whereby lack of full homology results in diminished binding, and lack of minimal microhomology (8-9 bp)(*24*), entirely abolished RecA filament binding.

Collectively, our supercoiling-calibrated assay recapitulates the established dependence of successful homology search on homology length and homologous sequence matching(*31–33*), while revealing that negative supercoiling permits efficient binding even by the shortest functional presynaptic filament.

### Homology search is determined by DNA supercoiling rather than DNA compaction

The prevailing model for RecA filament engagement with homologous DNA posits that efficient homology search requires a compacted, looped conformation of target DNA to enable ‘intersegmental transfer’(*15, 37*). In this model, a long, stiff filament rapidly samples multiple nearby DNA segments within the compacted DNA, captures a short microhomology region, and then aligns to adjacent sequences to complete binding. To determine if intersegmental transfer remains an important contributor to homology search in the presence of negative supercoiling, we conducted time-course binding assays across a wide range of DNA end-to-end extensions using a long, 500 nt, RecA filament. By varying flow speed during tethering, we generated surface-tethered molecules with end-to-end lengths spanning from ~5% (1.6 µm) to ~95% (13.5 µm) of our 42kb target’s contour length, while maintaining supercoiling density constant at σ = −0.05 (Fig. 2A-B, S4A). This range fully covers and extends the end-to-end distances explored in previous single-molecule studies(*15*).

**Figure 2.**
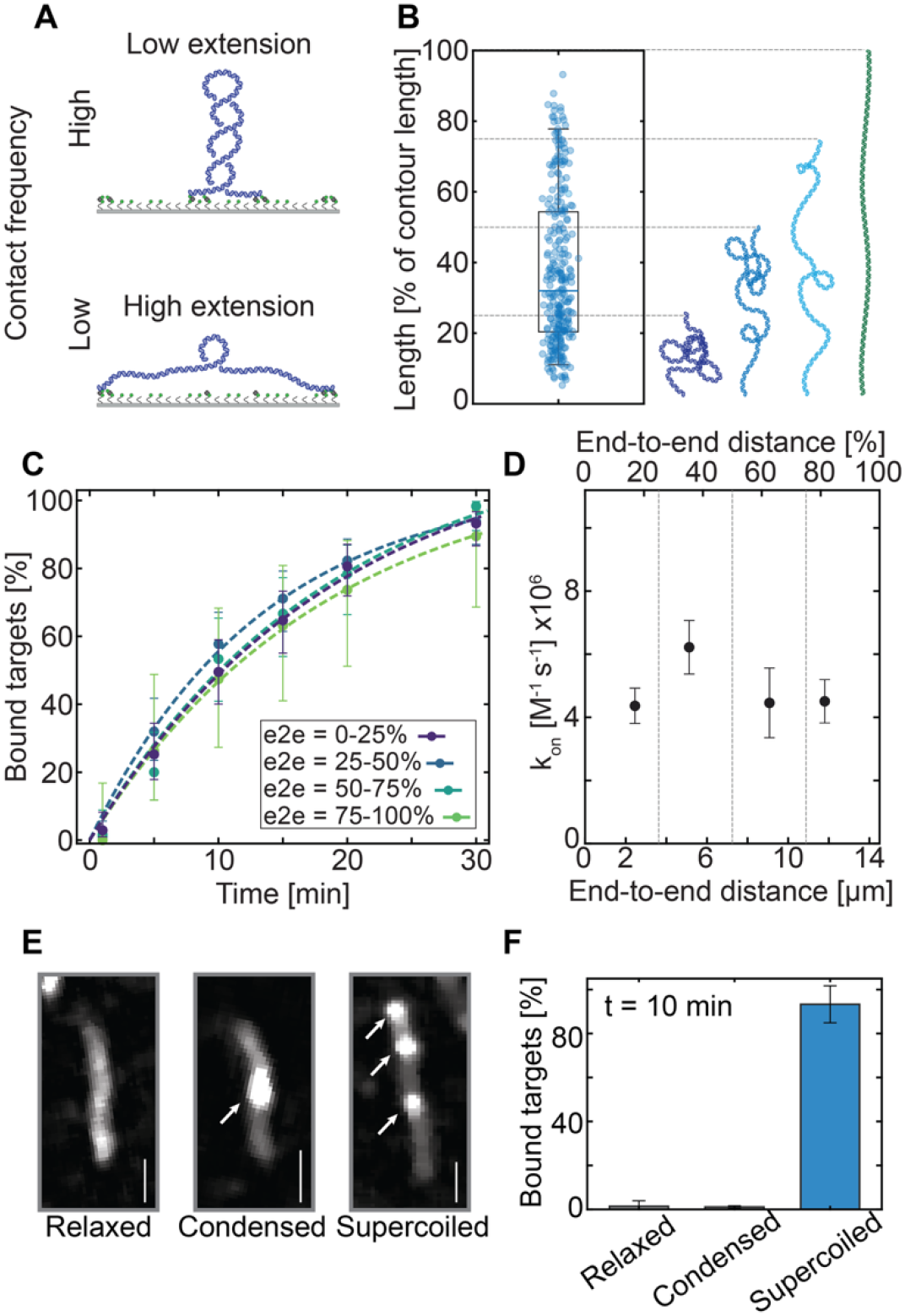
Productive homology search *in vitro* depends on DNA supercoiling rather than DNA compaction. A) Schematic representation of single-molecule assay. By varying the flow, dsDNA on the surface can be tethered at different end-to-end lengths, from fully compacted (~5% extension) to fully elongated (95% extension). **B)** Spread of the relative end-to-end extension in dsDNA target molecules on the surface during our experiments. Maximal coverage is from 4-98% of the contour length extension. Dashed lines represent the cut off of quartiles shown in panel C). n = 279 **C)** Timecourse experiments at different relative dsDNA extensions in the presence of 0.2 nM 500 nt RecA filaments (n_(0-25%)_ = 103, n_(25-50%)_ = 97, n_(50-75%)_ = 60, n_(75-100%)_ = 19). Dashed lines represent the fits of pseudo-first order kinetic binding model to the timepoint data. Error bars represent Wilson binomial confidence interval for each timepoint. **D)** Binding rates (k_on_) calculated by fitting the curves shown in C) with a pseudo-first order kinetic binding model. Error bars represent the standard error of the fitted pseudo-first-order rate constant. DNA length at 100% stretch corresponds to 14.5 µm (42 646 bp). **E)** Fluorescence images of the 42 kb dsDNA target molecule in relaxed (left), condensed (middle, 7.5% PEG 8000), and supercoiled (right, σ = −0.05) state. Scale bars = 2 µm. **F)** Fraction of bound dsDNA targets after a 10 min incubation with 0.2 nM 500 nt RecA filaments. Error bars represent standard deviation for each time point measurement (n = 3).

Collectively, binding dynamics were indistinguishable from those we observed at the intermediate (~50%) extensions used in our initial experiments (Fig.1, Fig. S4B), with nearly 100% of molecules achieving stable binding events. Segmenting molecules by their end-to-end extensions into quartiles revealed comparable binding kinetics (Fig. 2C), with apparent k_on_ values largely unchanged from the most compacted (<25% extension) to the most extended (>75% extension) molecules (Fig. 2D).

We observed the same complete independence of end-to-end distance for 90 nt RecA filaments which are too short to support meaningful intersegmental transfer. Notably, the 500 nt RecA filament, once assembled on ssDNA, spans ~255 nm (~0.51 nm/nt(*38*)). Using a worm-like chain model for relaxed B-form DNA, this RecA filament length enables effective sampling of microhomology segments separated by up to ~2800 bp along the DNA contour (Fig. S4C; see Methods). This is more than a 20-fold difference compared to ~100 bp for the 90 nt RecA filament. Regardless of the difference in sampling reach, binding rates showed no appreciable dependence on extension for both 90 nt, and 500 nt RecA filaments (Fig. 2D, S4D-G). Even when the DNA was stretched to more than 90% of its contour length, we still observed successful RecA filament binding within 30 minutes, indicating that DNA compaction is not required for efficient homology search (Fig. 2C, S4F, G).

To additionally control for compaction alone, we performed experiments on relaxed DNA (σ = 0) in the presence of a macromolecular crowding agent (Fig. 2E). We tethered the target DNA molecules at a low end-to-end distance (30 ± 7%, n = 85) to allow for mobility and spontaneous looping of the DNA, and then exposed them to 7.5% PEG which acts as a crowding agent. Crowding agent induced visible DNA compaction yet produced almost no binding (~1%), comparable to relaxed DNA without crowding (~2%) and in stark contrast to the ~93% bound fraction observed on negatively supercoiled DNA (Fig. 2F, S4H-I).

Taken together, our data show that DNA compaction does not impact homology search times and stable binding of RecA filaments. Rather, the presence of physiologically relevant level of negative supercoiling is the main determinant of successful homology binding.

### Negative supercoiling converts transient RecA sampling into committed homology binding

To determine which step of homology search is altered by DNA topology, we next followed homology search by individual RecA filaments in real time.

Stable RecA filament binding typically occurred in a single, abrupt step without detectable pre-binding diffusion along the target substrate. In instances of apparent RecA filament movement along the DNA, that movement coincided with large-scale plectoneme excursions, precluding unambiguous assignment of active search via one-dimensional diffusion proposed previously (D ≈ 8000 bp^2^/s or 9.25 x 10□^4^ µm^2^/s)(*39*) (Fig. 3A). Instead, observed motions likely arose from plectoneme dynamics that remain co-localized with the filament position (Fig. 3A-bottom).

**Figure 3.**
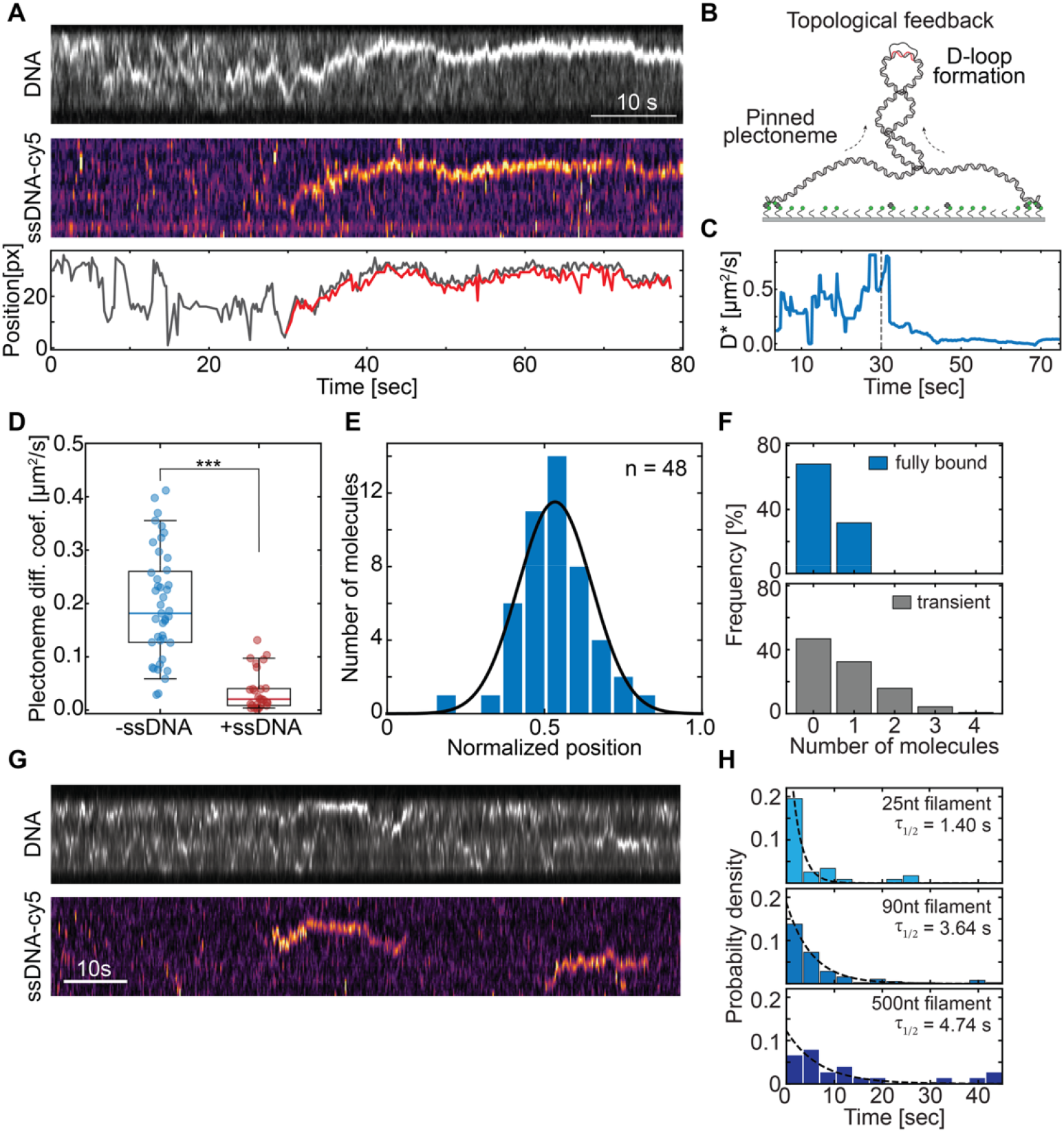
Real-time observation of homology search and plectoneme pinning. **A)** Top – Kymograph SxO-stained surface tethered dsDNA. Middle – kymograph for Cy5-labelled ssDNA-RecA presynaptic filament. Bottom – Overlay of peak fluorescence positions of dsDNA (gray) and ssDNA-RecA (red) channels. **B)** Schematic representation of pinned plectoneme at the D-loop position, shown in panel A. **C)** Time-resolved apparent diffusion coefficient (D*) of plectonemes on a dsDNA molecule shown in A), calculated as a rolling average over 6 s windows (30 frames). The dashed line indicates the time of ssDNA-RecA presynaptic filament binding and D-loop formation. **D)** Diffusion coefficients of plectonemes along individual dsDNA molecules, measured before (blue) and after (red) binding of the presynaptic filament. Each data point represents the average D from one DNA molecule (-ssDNA: n = 43, Mean ± SD = 0.195 ± 0.106 µm^2^/s; +ssDNA: n = 27, Mean ± SD = 0.037 ± 0.037 µm^2^/s). Boxplot whiskers indicate the 5-95th percentile range. Statistical significance: *** p < 0.001 (two-sided Mann– Whitney U test). **E)** Normalized binding positions of ssDNA-RecA presynaptic filaments along the dsDNA target (homology region placed at the center = 0.5). Black line - Gaussian fit to the data (µ = 0.53, σ = 0.12). n = 48 binding events. **F)** Two distinct modes of presynaptic filament interaction with dsDNA. Top (blue) - stable, prolonged binding with persistent plectoneme pinning (n = 44, 28% of total). Bottom (gray) - transient binding followed by dissociation (n = 113 events, 72% of total). x-axis shows the number of individual presynaptic filament binding events per dsDNA target molecule. See Fig. S5 for further details. **G)** Same as panel A, but for off-target transient binding events. **H)** Residence times of transient binding events for RecA filaments of different lengths (n_(25 nt)_ = 34, n_(90 nt)_ = 111, n_(500 nt)_ = 22). Black dashed lines represent exponential fit to data.

Strikingly, RecA filament binding persistently altered target DNA dynamics. Highly mobile plectonemes became pinned at the RecA filament binding site and co-localized with the filament throughout the experiment (Fig. 3A-B, Fig. S5A). Typical DNA plectoneme diffusion showed a quick transition from high diffusion coefficients (D ~ 0.1-0.5 µm^2^/s) to near-immobility (D ~ 0.02 µm^2^/s) upon RecA filament binding (Fig. 3C, D and S5B). These data indicate that stably bound RecA filaments form a mechanically or topologically distinct committed intermediate.

Stably bound RecA filaments were localized to the middle of the DNA target (Fig. 1E), well-aligned with the position of the homologous sequence. Our real-time data also revealed a different binding mode of RecA filaments; namely, transient binding which similarly altered DNA dynamics, but ultimately resulted in filament dissociation from the DNA target (Fig. 3F, G). In contrast to stably bound molecules, transient binding occurred at dispersed positions along the DNA target molecule, indicating off-target binding (Fig. S5C vs Fig. 3E) which suggests DNA sampling at regions of non-optimal homology(*40*). Longer RecA filaments experienced less frequent (Fig. S5D), albeit longer-lasting transient binding events (Fig. 3H). Transient binding was also present on relaxed DNA targets, however, with a shorter residence time (Fig. S5F) and lower binding frequency (Fig. S5G).

Importantly, we did not detect any transitions from transient to stable binding on relaxed DNA, in contrast to negatively supercoiled DNA, where stable complexes formed in 28% of RecA filament binding events (Fig. 3F, S5H). This suggests that negative supercoiling facilitates a transition from transient sampling to stable binding. Finally, despite short regions of microhomology along the target, filaments often bound stably at a single site per molecule, suggesting that the successfully detected homologous segment along with plectoneme pinning commits the DNA to the bound presynaptic filament via a yet unknown mechanism.

Overall, our real-time data demonstrate that RecA filaments are able to sample both supercoiled and relaxed DNA molecules, but negative supercoiling increases the sampling rate and permits the transition from transient sampling to stable homology binding.

### Preference towards negatively supercoiled DNA targets is broadly conserved in bacteria

RecA proteins are highly conserved across the bacterial kingdom and carry out HR in species with highly diverse chromosome architectures, lifestyles, and phylogenetic origins. We asked whether the strong preference of RecA filaments for negatively supercoiled targets is conserved across bacterial species, given that negative supercoiling levels may vary amongst different bacterial species(*41*). We therefore purified and compared RecA orthologs from Gram-negative - *E. coli* (presented throughout this study) and *C. crescentus* (CcRecA), and Gram-positive representatives - *Bacillus subtilis* (BsRecA), *Streptomyces venezuelae* (SvenRecA) and *Staphylococcus aureus* (SaRecA), spanning distant clades and ecological niches(*42*). All RecA orthologs maintained high sequence homology and showed comparable AlphaFold-predicted(*43*) RecA filament structure with right-handed architecture (Fig. S6A, B).

Under identical experimental conditions, using ATPγS-stabilized RecA filaments formed on 25 nt ssDNA, all orthologous RecA filaments showed efficient binding and strong discrimination in favor of negatively supercoiled DNA targets (Fig. 4), including SvenRecA which originates from a genus known for having linear chromosomes(*44*). BsRecA filaments displayed the highest binding to non-coiled DNA targets, however 97% of negatively supercoiled DNA was bound with BsRecA filaments within just 10 min, compared to 6% binding to relaxed DNA targets within the same time period. Notably, we were unable to achieve successful binding of BsRecA and SaRecA using 14 nt ssDNA, possibly due to the structural differences in the nucleation steps and stability. While orthologous RecA proteins showed differences in both binding kinetics (Fig. 4C) and ATPase rates (Fig. S6C-F), they all exhibited >100-fold increase in binding k_on_ to negatively supercoiled DNA targets compared to relaxed, non-coiled DNA targets.

**Figure 4.**
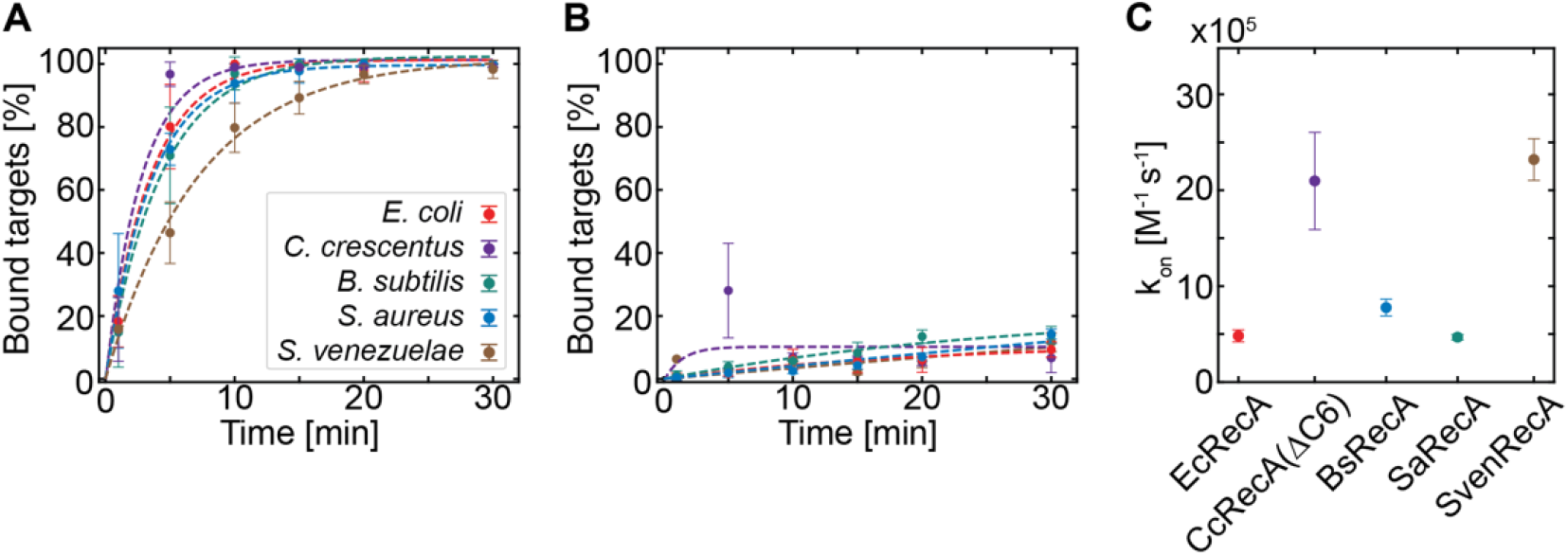
Extreme preference of RecA filaments for negatively supercoiled DNA is conserved across the bacteria. **A)** Timecourse experiments using 25 nt ssDNA as invasion strand and homologous RecA proteins from *Escherichia coli* (n = 334), *Caulobacter crescentus* (as CcRecA(ΔC6) mutant, n = 294), *Bacillus subtilis* (n = 322), *Staphylococcus aureus* (n = 293) and *Streptomyces venezuelae* (n = 302). Dashed lines represent the fits of pseudo-first order kinetic binding model to the timepoint data. Error bars represent standard deviation for each time point measurement (n = 3). **B)** Same as in A) but representing the nicked, torsionally unconstrained DNA surface tethered molecules. *E. coli* (n = 1403), *C. crescentus* (as CcRecA(ΔC6) mutant, n = 1307), *B. subtilis* (n = 1746), *S. aureus* (n = 1612) and *S. venezuelae* (n = 1457). **C)** Binding rates (k_on_) calculated by fitting the curves shown in A) with a pseudo-first order kinetic binding model, adjusted for concentration used in the experiments (EcRecA, SaRecA = 8nM, BsRecA = 5nM, CcRecA(ΔC6) = 3nM, SvenRecA = 1nM). Error bars represent the standard error of the fitted pseudo-first-order rate constant. Colors corresponding to panels A and B.

Overall, our data on five orthologs indicate that the extreme preference for negatively supercoiled substrates during homology search is a fundamental, widespread feature of bacterial RecA proteins.

### DNA supercoiling is essential for homology search in vivo

To understand how the topological dependence observed *in vitro* translates to the complex cellular environment, we observed RecA homology search in real-time in *Caulobacter crescentus* cells.

We used our previously established system in which an I-SceI site-specific double-strand break (DSB) is induced and RecA filament dynamics are tracked by fluorescence microscopy(*11, 45–47*). To allow repair from a homologous chromosomal locus, we halt cell division and ensure two fully segregated chromosomes in each cell. Low-level I-SceI expression generates a single double-strand break and results in the assembly of a RecA filament that performs homology search (Fig. 5A). We observe this in real-time using a fluorescent fusion of RecA-YFP(*11*).

**Figure 5.**
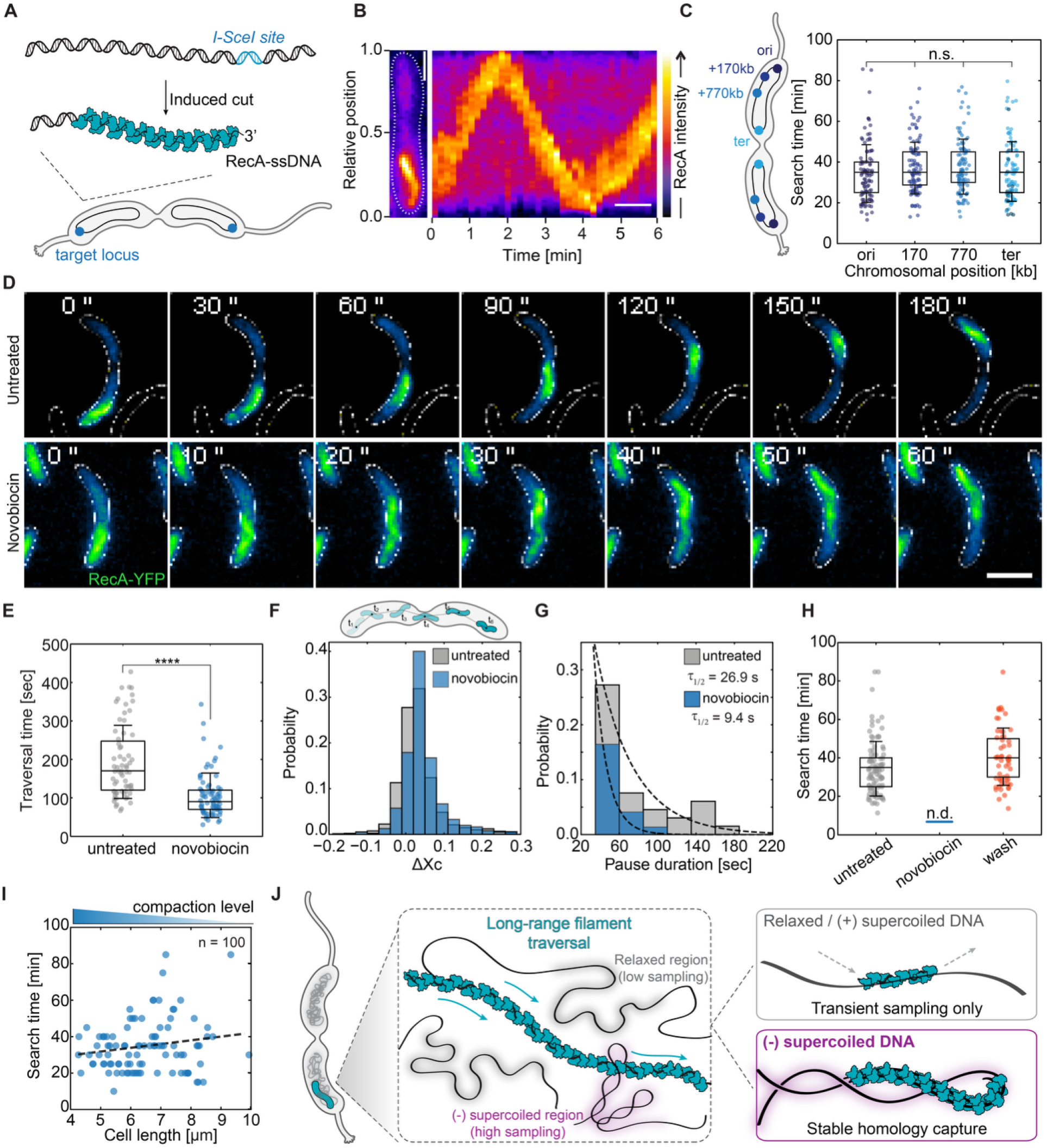
DNA supercoiling is essential for homology resolution *in vivo*. **A)** Schematic representation of the cellular loci targeted for double-stranded break (DSB). I-SceI restriction enzyme site is placed proximal to *Caulobacter ori* in this example. Vanillate induction of I-SceI expression leads to DSB and ssDNA formation. RecA proteins assemble on this nascent ssDNA to form a filament and search homology(*11, 45*). **B)** Kymograph of RecA-YFP position normalized to the cell length. Scale bar = 60 s. Left panel shows a single frame snapshot of RecA filament in a cell. Scale bar = 2 µm. **C)** Total homology search time at different DSB locations (30 kb/*ori*, 170 kb, 770 kb, 1958 kb/*ter*). n_*ori*_ = 100, n_*170kb*_ = 100, n_*770kb*_ = 100, n_*ter*_ = 100 Significance score: ns, using two-sided Mann-Whitney test. **D)** Timelapse tracking of RecA filament movement during homology search in the untreated cells (top) and cells treated with novobiocin (bottom) at 50 µg/ml for 30 minutes. Scale bar = 2 µm. Times, in seconds, indicated at each micrograph. **E)** RecA filament traversal time in untreated (gray, n = 66) and novobiocin treated (blue, n = 73) cells. Significance score: p < 0.0001, using two-sided Mann-Whitney test. **F)** Top - graphical depiction of per frame analysis. Centroid of RecA filament is tracked in each frame (10 s). Bottom - per frame displacement (ΔXc) distribution in untreated (gray) and novobiocin treated (blue) conditions, expressed as the distance relative to the cell length. **G)** RecA filament pause durations in untreated (gray) and novobiocin (blue) treated condition. Black dashed lines represent exponential decay fit to histogram data. **H)** Total homology search time of RecA filaments cut at *ori* location in different conditions. Gray - untreated cells (n = 100), blue/n.d - novobiocin treated cells (no observations of resolution, n = 0/100), orange - novobiocin treated cells with a subsequent wash out of novobiocin (40 ± 2 min, n = 58). Data for homology search time of RecA filament in untreated cells at *ori* has been reproduced from panel C for comparison. **I)** Total homology search time of RecA filaments relative to *C. crescentus* cell length. Linear fit correlation: R^2^ = 0.0293, n = 100. Black dashed line represents a linear regression fit to data **J)** Topology-centric homology search model. Mobile, extended RecA filament spans multiple regions and ensures high genome encounter and transient sampling, but higher sampling rate and productive homology pairing happens only in negatively supercoiled regions (purple halo).

As previously reported, induction of a double-strand break triggered the formation of a RecA filament that dynamically traversed the cell multiple times from pole to pole (Fig. 5B, S7A, Movie S2, S3). This allowed us to precisely quantify RecA filament speed, pole-to-pole traversal time, and total search time (search ends when the filament ceases traversal and rapidly disassembles). Interestingly, inducing the DSB at different locations spread across the chromosome (+0kb/*ori*, +70kb, +170kb, and +1950 kb/*ter*) showed identical traversal time and search time distributions across all positions (Fig. 5C, Fig. S7B-C). In all cases, the search time averaged ~37 min, despite large differences in physical distance between the cut and homologous locus (Fig. S7B). A subset of *ter*-proximal breaks resolved locally without large-scale filament movement, likely due to the proximity of the break- and repair-site (Fig. S7A, S7D). Nevertheless, the majority (83%) of *ter*-proximal breaks, involved long-distance traversal to complete the search. These baseline data establish that, once initiated, homology search proceeds via long-distance RecA filament traversal that is independent of the genomic position of break site or repair template.

We next asked how a loss of negative supercoiling would affect homology search. We thus exposed *C. crescentus* cells to novobiocin, a potent inhibitor of DNA gyrase(*48*), previously shown to abrogate negative supercoiling while preserving global chromosome organization(*49*). Accordingly, supercoiled plasmid species were no longer detected in *C. crescentus* cells after 30 min of treatment (Fig. S8A, B). In cells with comparable lengths (Fig. S8C), novobiocin treatment did not impair RecA filament formation at the break site (Movie S4-S5), yet the dynamics of traversal were markedly changed (Fig. 5D). Pole-to-pole traversal time reduced by ~45 % (106 ± 7 s versus 193 ± 12 s in untreated cells), showing that RecA filaments traversed the cells faster in the absence of supercoiling (Fig. 5E). Frame-to-frame displacement of the RecA filament centroid in the absence of novobiocin showed a positively skewed distribution with a prominent peak at 0, indicating that RecA filaments spend a substantial fraction of time in apparent pauses during traversal, interspersed with larger jumps that drive net movement (Fig. 5F, Fig. S8D). Consistent with an increased velocity, in novobiocin-treated cells this distribution shifted: the peak moved to positive values, reflecting more continuous, progressive movement toward the opposite pole with fewer and shorter pauses (Fig. 5F, S8E). Direct quantification of the pausing events revealed close to ~3-fold reduction in pause duration in novobiocin-treated cells compared to untreated controls (Fig. 5G).

Despite the accelerated and more continuous search by RecA filaments, homology resolution - i.e., finding the matching sequence, was entirely abolished. In untreated cells RecA filaments located and resolved at the homologous locus within ~37 min when a break was induced at *ori* (Fig. 5H). To our surprise, in novobiocin-treated cells we did not observe a single homology resolution event (0/100 cells) across a two-hour imaging window despite continuous pole-to-pole traversal by RecA filaments (Fig. 5H). These data show that in the absence of negative supercoiling RecA filaments moved faster, paused less, yet remained unable to achieve homology recognition. Importantly, the effects of novobiocin were fully reversible. Applying identical novobiocin treatment, and then washing out the drug restored both normal dynamics and homology resolution upon break induction (Fig. 5H). We observed the same, perpetual RecA traversal and a complete block of homology resolution when the break was induced at the *ter* locus as well, demonstrating that supercoiling was important for homology search irrespective of the location of the break-site or homologous template (Fig. S8F).

Finally, we tested whether novobiocin treatment impaired resolution indirectly by altering chromosome compaction rather than supercoiling. In *C. crescentus*, cell length is a direct proxy for nucleoid compaction levels(*49*). Longer cells have less condensed chromosomes, shown by proportionally greater distances between chromosomal loci (Fig. S8G-H). If compaction were a primary determinant of search efficiency, search time should increase with cell length. Instead, search times did not show any correlation with cell length across a wide range of cell sizes (R^2^ < 0.02, Fig. 5I). We also did not observe a correlation between traversal times and cell lengths (R^2^ < 0.02, Fig. S8I), consistent with the idea that RecA filament mobility is primarily constrained by the DNA which it is scanning and not the cell boundaries. This mirrors our *in vitro* observations where DNA supercoiling, rather than DNA compaction, dictates productive homology pairing. Together, our data establish that DNA supercoiling is the key determinant of a productive homology search.

## Discussion

Our data establish that the topological state of DNA is not an accessory feature of the genome, but an essential gatekeeper of homology search. Decades of previous research provided sparse hints that negatively supercoiled DNA is required for homology search, notably in magnetic tweezers experiments that required negative supercoiling to observe RecA strand exchange (*23, 50*) and in biochemical assays on plasmid DNA that measured topological changes after D-loop formation (*21*). Here we provide the direct causal link between DNA topology and homology search. By directly controlling DNA supercoiling *in vitro* and perturbing chromosomal supercoiling during homology search *in vivo*, we show that DNA supercoiling is the determining physical property that allows successful target capture. In this way, our work resolves a long-standing problem in homologous recombination: not simply how RecA filament samples the genome, but also how those encounters become productively committed to repair.

On a molecular level, a RecA filament samples the target DNA molecule through transient encounters that do not progress to stable pairing unless negative supercoiling is present. Negative supercoiling likely lowers the energetic barrier for base flipping and target DNA opening observed in structural models of the RecA nucleoprotein complex(*14, 34*). Our data firmly support this. *In vitro*, RecA filaments sampled both relaxed and negatively supercoiled substrates (Fig. 3, Fig. S5), but negative supercoiling supported more frequent sampling and, uniquely, their progression to stable homology binding. This effect carried over seamlessly *in vivo*. In cells, we observed RecA filaments pausing at sites distant from homologous targets, which likely represent microhomology interrogation. Loss of negative supercoiling did not prevent RecA filament formation or long-range traversal. Instead, RecA filaments moved faster, paused less, and yet completely failed to resolve homology. These observations answer how homologous targets are located amid a vast excess of non-homologous sequences: genome-wide negative supercoiling increases both the effective sampling rate and the probability of stable binding once homology is encountered.

Our topology-centric view reframes the question of homology search by positioning genome-wide supercoiling as the main selectivity filter downstream of filament mobility or extension. ‘Reduced-dimensionality search’ enabled by extended (*10*) or mobile (*11*) RecA filaments can accelerate RecA filament encounters with the homologous target, yet the final success is determined by the presence of DNA supercoiling. In contrast, ‘intersegmental transfer’ within compacted DNA (*15, 37*) is not required for efficient homology search and capture. This was shown both by our *in vitro* and *in vivo* experiments where the degree of DNA compaction had no effect on overall search times (Fig. 2, Fig. 5). Instead, we offer a new framework: RecA filaments sample the genome for homology through dynamic motion and filament extension (*10, 11*), while productive pairing occurs only when the local supercoiling landscape permits strand engagement and subsequent stabilization of the bound RecA filament (Fig. 5J). In this framework, mobility solves the encounter problem, whereas DNA topology solves the commitment problem.

These findings carry broad implications for genome maintenance. Replication (*2, 51*) and transcription (*52, 53*) strongly induce negative supercoiling upstream of their respective machineries, thereby continuously generating local supercoiling gradients across the chromosome. Such gradients could actively guide RecA filaments toward regions of appropriate topology, thereby directing repair to recently replicated and transcribed loci where negative supercoiling is enriched. In this way, the cell harnesses its intrinsic topological landscape as a powerful guide for repair, increasing the efficiency of homology search without relying on sequence information alone.

Given the high functional conservation of mechanisms such as homologous recombination, DNA supercoiling, DNA replication and transcription across bacteria, a topology-centric homology search is likely an ancient mechanism that couples genome topology to DNA repair. Our data demonstrate that the preference for negatively supercoiled substrates is a widespread feature of bacterial RecA proteins (Fig. 4). Whether analogous topological selectivity operates in eukaryotic recombination systems remains an open and intriguing question, especially given that SMC complexes, which guide homologous recombination in human cells (*54, 55*), also universally interact with-(*56–58*) and alter DNA supercoiling (*59–61*).

Overall, our results establish DNA supercoiling as a genome-wide selectivity filter that is essential for efficient and faithful homologous recombination, with direct and far-reaching implications for longstanding *in vitro* and *in vivo* recombination models.

## Materials and Methods

### Preparation of the DNA substrate for surface tethering

The surface-tethered linear DNA used in the single-molecule imaging assay is based on a previously reported 42 kb plasmid(*62, 63*), which is stably propagated in *E. coli* NEB 10-beta cells at 28°C. Additionally, a ~540 nt sequence that contains three chi sites and one I-SceI cleavage site(*10*) (Table S3) was introduced into the original plasmid at the unique NdeI site by isothermal assembly, generating pMT004 (pBS-chi3) (SE1019). Correct insertion of the sequence was confirmed by nanopore sequencing.

To prepare a linear fragment for surface tethering, the pMT004 plasmid was isolated from a 50 ml overnight culture in LB supplemented with 100 µg/ml ampicillin using a midiprep kit (Invitrogen). The non-target sequence construct derived from pBS-parS, described previously(*63*) (Fig. S3A-C), was prepared in the same way. Approximately 30 µg of plasmid DNA was treated with RecBCD enzyme (Exonuclease V, New England Biolabs) in the presence of 1 mM ATP at 37°C for 15 min, followed by enzyme inactivation at 70°C for 30 min. The recommended addition of 11 mM EDTA was omitted to avoid interference with subsequent reaction steps. Residual circular plasmid was then digested with NotI-HF (New England Biolabs) for 3 h at 37°C, followed by enzyme inactivation at 80°C for 20 min. Biotinylated end fragments (handles) were generated either with 5′-biotinylated primers (T43/T44) or, for most of the work presented here, with standard primers of identical sequence (T21/T22) (see Table S1). For internal biotin incorporation using the T21/T22 primer pair, PCR was performed with Taq polymerase and pMT004 as template in the presence of 1 mM dATP, dGTP, and dCTP, 0.75 mM dTTP, and 0.25 mM biotin-11-dUTP (Lumiprobe, Cat. #1553). Amplification of this 1282 bp product resulted in stochastic incorporation of biotinylated deoxynucleotides into the PCR fragment. The biotinylated PCR product was digested with NotI-HF using the same procedure as for pMT004, consisting of 2 h at 37°C followed by 20 min at 80°C, yielding two ~600 bp DNA handles.

The DNA handles were then added to the cut pMT004 at an approximately 15:1 molar ratio together with T4 DNA ligase (New England Biolabs) and 1 mM ATP, and the ligation reaction was incubated at 16°C for 16 h. To remove excess biotinylated handles, the ligation mixture was purified on a HiPrep™ Sephacryl™ S-500 16/60 gel filtration column (Cytiva) equilibrated in TE buffer supplemented with 150 mM NaCl. The sample was run at 0.2 ml/min, and fractions were collected manually into 1.5 ml Eppendorf tubes. All collected fractions were stored at 4°C until use, as freeze-thaw cycles tended to compromise DNA integrity. The final mixture contained both coilable and non-coilable DNA molecules, allowing direct comparison in the single-molecule assays.

### Purification of E. coli SSB protein

Recombinant *E. coli* SSB was purified from Ros*eff*a2(DE3) pLysS pET21-his10-sumo-ssb (SE204). We inoculated LB medium supplemented with ampicillin (70 µg/ml) and chloramphenicol (21 µg/ml) with an overnight culture, expanded the culture the following day, and induced protein expression with 1 mM isopropyl β-D-1-thiogalactopyranoside (IPTG) at OD600 = 0.65. After induction, the culture was transferred to 30°C and incubated with vigorous aeration for ~5.5 hours before harvesting the cells. The pellet was resuspended in lysis buffer [50 mM Tris-HCl pH 7.5, 200 mM NaCl, 1 mM EDTA, 5 mM β-ME, 10% sucrose, 0.1 mM PMSF, Benzonase, and cOmplete™ Protease inhibitor (Roche)] at 5 ml/g pellet, homogenized, supplemented with lysozyme to 1 mg/ml, incubated on ice for 20 min, and lysed using a microfluidizer (LM10, Microfluidics). Cell debris was removed by centrifugation at ~100 000 g for 40 min at 4°C, and the supernatant was adjusted to 30 mM imidazole and incubated with Ni-NTA resin for 30 min at 4°C. The resin was washed sequentially with high-salt washing buffer [50 mM Tris-HCl pH 8.3, 1 M NaCl, 50 mM imidazole, 5 mM β-ME, 10% glycerol] and low-salt washing buffer [50 mM Tris-HCl pH 8.3, 100 mM NaCl, 50 mM imidazole, 5 mM β-ME, 10% glycerol]. His_10_-SUMO-SSB was eluted in elution buffer [50 mM Tris-HCl pH 8.3, 250 mM NaCl, 200 mM imidazole, 5 mM β-ME, 10% glycerol]. His10-SUMO was cleaved off by His_6_-Ulp1, which was added to the preparation during the overnight dialysis against dialysis buffer [20 mM Tris-HCl pH 8.3, 500 mM NaCl, 1 mM EDTA, 2 mM β-ME, 10% glycerol]. After removing the cleaved His_10_-SUMO tag and His_6_-Ulp1 with Ni-NTA, the fraction containing SSB was dialyzed overnight against 1.5l of the dialysis buffer, and concentrated using a concentrator (MWCO 10 kDa). The concentrated preparation was diluted to reduce the concentration of NaCl to ~25 mM, and loaded onto a HiTrap Heparin HP column (Cytiva). SSB was eluted using a 0 - 1000 mM NaCl gradient in Hep A buffer [20 mM Tris-HCl pH 8.3, 1 mM EDTA, 5 mM β-ME, and 20% glycerol]. Elution fractions that contain SSB were pooled, concentrated, and further purified by size-exclusion chromatography (SEC) on a Superdex 75 10/300 GL column (Cytiva) equilibrated in SEC buffer [20 mM Tris-HCl pH 8.3, 500 mM NaCl, 1 mM EDTA, 5 mM β-ME, 20% glycerol]. Fractions that contain SSB were pooled and concentrated, flash frozen in liquid nitrogen, stored at −80°C. The concentration was determined based on the absorbance at 280 nm.

### Purification of E. coli RecA

Recombinant *E. coli* RecA was purified using multiple rounds of polyethylenimine and ammonium sulfate precipitation as previously described by Cox et al.(*64*), using an expression vector pDuet1-RecA (pMT007). We constructed this vector using NcoI-cut pETDuet1 plasmid (MilliporeSigma (Novagen)) and *recA* gene extracted by PCR from the *E. coli* genome using primers SC2080/SC2081.

First, we inoculated 50 ml of LB medium supplemented with 100µg/ml of ampicillin (LB-Amp) with Ros*eff*a2 pLysS cells carrying pETDuet-recA (pMT007) plasmid and grew this overnight at 37°C with continuous shaking at 200 rpm. The following day, we inoculated 4 L of LB-Amp using the overnight culture to a final dilution of 1:100 (i.e., 10 ml : 1 L). Large culture was grown to mid-exponential phase (OD = 0.7) and then moved to 4°C to slow down the growth for 20 min. We induced RecA expression by adding 1 mM IPTG and transferred the large culture to 16°C, for further expression at 16°C for 20 h. Cells were harvested at 7000 g for 15 min, resuspended in lysis buffer [50 mM Tris-HCl pH 7.5, 1 mM EDTA, 10 mM β-ME, 15% sucrose] at 1 g/ml, lysed using a microfluidizer, and clarified by centrifugation at 150 000 g for 30 min. Polyethylenimine was added to the supernatant to a final concentration of 0.5%, and after 30 min incubation at 4°C the precipitate was collected by centrifugation at 14 000 g for 15 min. The pellet was extracted sequentially with R buffer [20 mM Tris-HCl pH 7.5, 10 mM β-ME, 0.1 mM EDTA, 10% glycerol] containing 150 mM and then 300 mM ammonium sulfate, each for 30 min, followed by centrifugation at 13 000 g for 15 min. Solid ammonium sulfate was then added to the recovered supernatant to 0.28 g/ml, and after 20 min stirring the precipitated protein was collected by centrifugation at 26 000 g for 20 min. The pellet was resuspended in dialysis buffer [50 mM Tris-HCl pH 7.5, 1 mM EDTA, 200 mM NaCl, 10 mM β-ME] and dialyzed three times against the same buffer (2 h, 18 h, 2 h). The dialyzed sample was diluted to ~70 mM NaCl in HA buffer [50 mM Tris-HCl pH 7.5, 1 mM EDTA, 10 mM β-ME], loaded onto a HiTrap Heparin HP column pre-equilibrated in HA buffer containing 10 mM NaCl, and eluted with a 10-1000 mM NaCl gradient over 40 ml. RecA eluted at ~400-450 mM NaCl (conductance ~23-27 mS/cm). Peak fractions were pooled, dialyzed twice against RecA storage buffer [20 mM Tris-HCl pH 7.5, 200 mM NaCl, 0.1 mM EDTA, 1 mM DTT, 5% glycerol], concentrated, snap frozen in liquid nitrogen, and stored at −80°C in 10 µl aliquots.

### Labeling of E. coli RecA protein

For experiments with unlabelled 500 nt ssDNA (MT003), we used Cy5.5-labelled RecA protein. Purified RecA was labelled at the N-terminal amine with Sulfo-Cyanine5.5 NHS ester as previously described(*65*) with the following modifications. For labeling, 200 µl of highly concentrated RecA was diluted 12.5-fold in phosphate buffer [NaH2PO4/Na2HPO4 pH 7.0, 500 mM NaCl] to a total volume of 2.5 ml and loaded onto a PD-10 Sephadex G-25 column (Cytiva) pre-equilibrated with the same buffer. The eluted RecA fraction, now free of storage-buffer components, was concentrated to 250 µl using a 10 kDa MWCO Amicon® Ultra centrifugal filter (MilliporeSigma). Sulfo-Cyanine5.5 NHS ester (Lumiprobe) was then added in 10-fold molar excess, and the reaction was incubated in the dark for 15 min at room temperature followed by 3 h on ice. To remove unreacted dye and exchange buffers, the labeling reaction was diluted in RecA storage buffer, thereby quenching excess Cy5.5 NHS ester, and passed through a PD-10 Sephadex G-25 column pre-equilibrated with RecA storage buffer containing 150 mM NaCl. Residual unreacted dye was removed by repeated rounds of dilution and concentration using a 10 kDa MWCO Amicon® Ultra centrifugal filter. In each round, the protein was concentrated to 400 µl and then diluted 10-fold in storage buffer to further reduce residual dye carried over from the column. This procedure was repeated three times, after which the sample was concentrated to 200 µl, snap frozen in liquid nitrogen, and stored at −80°C in 10 µl aliquots until use. The removal of excess dye was assessed by 5-15% SDS-PAGE gel and imaging the gel for fluorescence of Cy5.5 on a bioimager (Ex = 645nm, Amersham™ Typhoon™). We determined the protein labeling efficiency at 37% by using extinction coefficients of 21 430 M^−1^cm^−1^ and 211 000 M^−1^cm^−1^ for RecA and Sulfo Cy5-NHS, respectively, with a correction factor at 260 nm wavelength of CF_260_ = 0.09 as recommended by Lumiprobe.

### Purification of homologous RecA proteins (BsRecA, CcRecA, SaRecA, SvenRecA)

For expression of homologous RecA proteins, we constructed pETDuet1-based expression vectors by assembling NcoI-linearized pETDuet1 with synthetic gene fragments carrying codon-optimized *recA* orthologs using a 1:4 vector:insert HiFi reaction (see Table S4) and transformed the products into ECOS™ Sonic™ competent cells lacking endogenous *recA* gene.

Overnight cultures grown in 60 ml LB-Amp were used to inoculate 6 L LB-Amp at 1:100 dilution. Cultures were grown to OD600 = 0.7, cooled for 20 min at 4°C, induced with 0.4 mM IPTG, and expressed at 16°C for 20 h. Cells were harvested at 8000 g for 20 min, resuspended in lysis buffer [50 mM Tris-HCl pH 7.5, 1 mM EDTA, 10 mM β-ME, 10% sucrose, 0.1 mM PMSF, cOmplete™ Protease inhibitor (Roche)] at 5 ml/g pellet, supplemented with lysozyme to 1 mg/ml and incubated on ice for 20 min. Next, the cells were lysed by sonication (2s ON, 2s OFF, 40% amplitude for 2 min, repeated twice), and centrifuged at 35 000 g (Ti45, Beckman Ultracentrifuge). The supernatant was treated with 0.6% polyethylenimine for 20 min at 4°C, and the pellet was extracted sequentially with R100 buffer [40 mM Tris-HCl pH 7.5, 100 mM NaCl, 0.1 mM EDTA, 5 mM β-ME, 10% glycerol, cOmplete™ protease inhibitor] and R900 buffer [40 mM Tris-HCl pH 7.5, 900 mM NaCl, 0.1 mM EDTA, 5 mM β-ME, 10% glycerol, cOmplete™ protease inhibitor]. The R900 extract was diluted to 500 mM NaCl, precipitated with 0.22 g/ml ammonium sulfate, and the recovered pellet was resuspended in high-salt dialysis buffer [25 mM Tris-HCl pH 7.5, 500 mM NaCl, 1 mM EDTA, 5 mM β-ME, 30% glycerol] and dialyzed twice against 1.5 L of the same buffer (2 h and 16 h). The dialyzed sample was diluted to 70 mM NaCl in HA buffer [25 mM Tris-HCl pH 7.5, 10 mM NaCl, 1 mM EDTA, 5 mM β-ME, 10% glycerol], loaded onto a 5 ml HiTrap Heparin HP column, and eluted with a 20-1000 mM NaCl gradient.

Peak fractions were diluted to 150 mM NaCl in Blue A buffer [40 mM HEPES pH 7.5, 100 mM NaCl, 1 mM EDTA, 1 mM DTT, 5% glycerol], loaded onto a 5 ml HiTrap Blue HP column, and eluted with a 0-100% gradient into Blue B buffer [40 mM HEPES pH 7.5, 2000 mM NaSCN, 1 mM EDTA, 1 mM DTT, 5% glycerol]. Peak fractions were dialyzed overnight against HA buffer containing 300 mM NaCl, diluted to 50 mM NaCl, and further purified on a 1 ml HiTrap Q HP column using a 0-1000 mM NaCl gradient. Final peak fractions were dialyzed against storage buffer [20 mM Tris-HCl pH 7.5, 300 mM NaCl, 0.1 mM DTT, 10% glycerol]. SvenRecA, CcRecA, and CcRecA(ΔC6) lost ATPase activity after HiTrap Blue HP purification, and for these variants the HiTrap Blue HP column was omitted. Because full-length CcRecA did not show productive binding in our single-molecule assays despite retaining ATPase activity, we used a CcRecA(ΔC6) mutant lacking the final six C-terminal residues, generated by PCR linearization of pMT011 with phosphorylated primers T46/T49 followed by gel purification and ligation. SvenRecA, CcRecA, and CcRecA(ΔC6) were stored in higher-salt storage buffer [20 mM Tris-HCl pH 7.5, 500 mM NaCl, 0.1 mM DTT, 10% glycerol] because of their high aggregation propensity.

Proteins were concentrated the following day using 10 kDa MWCO Amicon® Ultra filters to a maximum of 150 µM, and concentrations were determined using extinction coefficients of 18 910 M^−1^cm^−1^, 17 420 M^−1^cm^−1^, 15 930 M^−1^cm^−1^, and 14 440 M^−1^cm^−1^ for SaRecA (*S. aureus*), BsRecA (*B. subtilis*) and CcRecA (*C. crescentus*), SvenRecA (*S. venezuelae*) respectively.

### Determination of ATP hydrolysis activity in RecA orthologs

To assess ATP hydrolysis activities of our purified RecA orthologs we performed real-time measurements of ATP hydrolysis and inorganic phosphate release using EnzChek® Phosphate Assay Kit (E-6646, ThermoFisher Scientific). All reactions were set up in identical concentrations and buffers for direct comparability. The final RecA binding reaction contained [20 mM Tris-OAc pH 7.5, 10 mM Mg(OAc)_2_, 20 mM potassium glutamate, 1 mM DTT, 10% (v/v) glycerol, 1U/ml purine nucleoside phosphorylase (PNP), 0.2 mM 2-amino-6-mercapto-7-methyl-purine riboside (MESG)], with an addition of 0.1 mM ssDNA (MT18, Table S2) and 0.5 µM RecA homolog. The reactions were split into separate wells of a Corning® 96 well plate (MiliporeSigma), to the final volume of 100 µl following the recommended EnzChek® protocol. The reactions were started by adding 0 - 1 mM ATP (See Fig. S6), and incubated at 37°C for 90 min, with continuous monitoring of absorbance signal at 360 nm, appropriate for PNP and MESG reaction product. All raw data were converted from OD_360_ to inorganic phosphate concentration (in µM), corrected for inorganic phosphate background, and normalized using absorbance values at 0 mM ATP, before calculating Michaelis-Menten kinetics and catalytic rates (Fig. S6).

### Surface passivation and preparation

Single-molecule experiments were performed using custom-made flow cells, assembled by connecting surface-passivated glass slide and glass coverslip using double-sided tape to segment individual lanes. Cleaning of surfaces was performed as described in detail by Chandranoss *et al*.(*66*), except that glass slides were used in all experiments. To remove non-specific particles, the surfaces were silanized in a solution of 10% (v/v) (3-aminopropyl)trimethoxysilane (APTMS; Sigma Aldrich) and 5% (v/v) acetic acid in methanol for 25 min, washed extensively with methanol, and dried at 70°C for 60 min. PEG passivation was then carried out using a mixture of 70 mg/ml methoxy PEG succinimidyl valerate (5,000 Da) and 15 mg/ml biotinylated PEG succinimidyl valerate (5,000 Da) in collapsed PEG buffer [0.55 M K2SO4, 0.1 M Na2CO3/NaHCO3 pH 9.6], which provides ‘cloudy’ PEG conditions and promotes full surface coverage(*67*). This solution was incubated with the slides for 24 h at 4°C, after which the surfaces were washed extensively with water to remove unbound PEG. Two additional rounds of PEGylation were then performed for 24 h each at 4°C using 10 mg/ml MS(PEG)4 methyl-PEG-NHS ester (ThermoFisher Scientific) and 0.5 mg/ml biotinylated PEG succinimidyl valerate (5,000 Da) in collapsed PEG buffer, with extensive washing with water between rounds to ensure complete passivation. On the day of the experiment, the surfaces were treated with 0.5 mg/ml UltraPure™ BSA (ThermoFisher Scientific) for 30 min in imaging buffer A [40 mM Tris-acetate pH 7.5, 60 mM potassium glutamate (K-glut), 10 mM magnesium acetate (Mg(OAc)_2_), 1 mM DTT].

To immobilize our 42 kb DNA, we flowed ~10 pM of biotinylated DNA molecules at flow rates of 6 µl/min in imaging buffer B [40 mM Tris-Acetate pH 7.5, 60 mM potassium glutamate (K-glut), 10 mM magnesium acetate (Mg(OAc)_2_), 1 mM DTT, 0.25 mg/ml UltraPure™ BSA, 250 nM SYTOX Orange (SxO)] until the surface coverage was satisfactory. In experiments in which end-to-end distance was varied, the flow rate was adjusted between 1 and 30 µl/min to achieve the desired DNA extension. The chamber was then washed with imaging buffer B lacking DNA to remove unbound molecules. DNA supercoiling was introduced into surface-tethered molecules by varying the SxO concentration before and after the DNA became topologically constrained, as described previously(*25*). For positive supercoiling, DNA was first tethered in imaging buffer B lacking SxO and then washed with imaging buffer B containing 40 nM SxO. For negative supercoiling, DNA was initially tethered in the presence of high SxO concentrations (100-1800 nM) and then washed with imaging buffer B containing 40 nM SxO. The presence of supercoiling was confirmed by the appearance of rapidly diffusing DNA plectonemes in surface-tethered molecules (e.g., Fig. S1D, Movie S1).

### RecA presynaptic filament assembly and binding

Real-time imaging of RecA-ssDNA binding to dsDNA targets was performed by flowing pre-assembled presynaptic filaments into the flow cell. Presynaptic filaments were assembled by mixing 0.5 µM Cy5-labelled ssDNA with RecA in RecA folding buffer [50 mM Tris-OAc pH 7.5, 10 mM Mg(OAc)_2_, 1 mM DTT, 1 mM ATPγS]. The RecA concentration was adjusted to maintain a constant stoichiometry of one RecA monomer per 2.9 nucleotides of ssDNA (for example, ~15.5 µM RecA for 90 nt ssDNA). Carryover salt from the RecA storage buffer was kept below 20 mM NaCl. For short ssDNA substrates (7-25 nt), in-house purified RecA and commercial RecA (New England Biolabs) were used interchangeably, as no difference in binding was observed. For unlabelled 500 nt ssDNA presynaptic filaments, unlabelled RecA and Cy5-RecA were used at a 10:1 molar ratio, with a total stoichiometry of one RecA monomer per 2.5 nucleotides of ssDNA. In experiments with RecA orthologs, the folding buffer was supplemented with 20 mM K-glut and 20% (v/v) glycerol to reduce aggregation and improve presynaptic filament assembly. Assembly reactions were incubated at 37°C for 1 min for E. coli RecA and for 3 min for homologous RecA proteins (BsRecA, SaRecA, CcRecA, and SvenRecA). Assembly reactions with 500 nt ssDNA were incubated for 5 min at 37°C with occasional mixing to ensure complete RecA binding.

The presynaptic filaments were then introduced at the desired concentrations (0.2-10 nM) in imaging buffer C [40 mM Tris-acetate pH 7.5, 60 mM K-glut, 10 mM Mg(OAc)_2_, 1 mM DTT, 0.5 mM ATPγS, 0.25 mg/ml UltraPure BSA, 40 nM SxO, 2 mM protocatechuic acid (PCA), 50 nM PCA dehydrogenase (PCD), 1 mM Trolox (6-hydroxy-2,5,7,8-tetramethylchroman-2-carboxylic acid)], and imaging was started immediately. Because CcRecA(ΔC6) and SvenRecA showed strong aggregation and non-specific binding as large clusters on both supercoiled and nicked DNA at high concentrations, the concentrations used for these variants were reduced from 5-10 nM, as used for EcRecA, BsRecA, and SaRecA, to 1 nM for SvenRecA and 3 nM for CcRecA(ΔC6).

For experiments performed under molecular crowding conditions (Fig. 2E, Fig. S4H), PEG 8000 (Thermo Fisher Scientific) was added to a modified imaging buffer C lacking BSA [40 mM Tris-acetate pH 7.5, 60 mM K-glut, 10 mM Mg(OAc)_2_, 1 mM DTT, 0.5 mM ATPγS, 40 nM SxO, 2 mM PCA, 50 nM PCD, 1 mM Trolox]. The flow cell was first equilibrated with buffer containing 5% (w/v) PEG 8000 at 2 µl/min for 10 min. Buffer containing 7.5% (w/v) PEG 8000 together with 0.2-0.5 nM RecA filament (500 nt) was then introduced, and imaging was started immediately.

### Single-molecule fluorescence imaging

We carried out real-time imaging using a home-built objective-type TIRF microscope with alternating excitation of 561 nm (0.4 mW) and 647 nm (10 mW) lasers in Highly Inclined and Laminated Optical sheet (HiLo) mode, as described in details in Ref. (*68*), to image SxO-stained DNA and Cy5-labelled ssDNA, respectively. All images were acquired with an sCMOS camera (ORCA-Quest, Hamamatsu) at an exposure time of 100 ms per color, thus 200 ms frame rate, with a 60x objective (Olympus PlanAPo 60X, 1.45 NA).

### Real-time RecA-ssDNA image analysis

Areas with single DNA molecules were cropped from the original raw images and analyzed separately using a previously described python analysis software by Pradhan et al.(*69*). For the analysis, cropped images were smoothened using a median filter (window size = 10 pixel - px), and background subtracted with the white_tophat operation in python module - *scipy*. The ends of each DNA molecule were manually marked. Kymographs were generated by summing the total fluorescence intensity across 14 pixels perpendicular to the DNA axis and stacking these values over time (Fig. 3). Peaks of high intensity in both the SxO-labelled DNA and Cy5-labelled ssDNA channels were identified using *scipy* and merged into tracks using a 10-frame time window and a 10 px lateral search window. To calculate apparent diffusion coefficients, peaks were linked into continuous trajectories using *trackpy.link_df* with a search range of 5 pixels and a memory of 5 frames, and only tracks spanning at least 10 frames were retained.

For each movie, trajectories were cropped to a symmetric window of ±75 frames centered on the manually annotated ssDNA binding time, separately for the pre-binding and post-binding intervals. Within each window, displacement was analyzed under a one-dimensional diffusion model as a function of lag time, τ. For each value of τ, all pairs of peak positions separated by exactly τ frames were identified, squared displacements were calculated, and the values were averaged to obtain MSD(τ), including cases with non-consecutive frames caused by detection gaps. A linear regression was then applied to the MSD values for τ = 2-10 frames (0.2-2.0 s) to obtain the diffusion coefficient, *D*, in px^2^/frame. Diffusion coefficients were converted to physical units (µm^2^/s) using a pixel size of 0.154 µm/pixel and a frame interval of 0.2 s (200 ms). Per-track diffusion coefficients were averaged to obtain one pre-binding and one post-binding value for each movie, and these values were then compiled across all analyzed molecules for the statistical comparison shown in Fig. 3D.

### Fitting experimental data and analysis

We fitted the time-course data for the percentage of negatively supercoiled DNA molecules bound by RecA-ssDNA using a pseudo-first-order kinetic model, assuming a large excess of presynaptic filaments over the tethered DNA targets on the surface. Raw percentage data was grouped by time points and computed for mean and standard deviation for each time point. Exponential growth model was fit to the mean values:

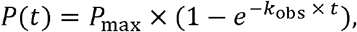

where *P*(*t*) is the percentage at time *t, P*_max_ is the maximum percentage bound, and *k*_obs_ is the observed pseudo-first-order rate constant (in min^−1^). Half-time *t*_1/2_ was derived as ln(2)/*k* _obs_ for cases where *k*_obs_ > 0. In cases where the fitted *P*_max_ was ≤ 5%, *k*_obs_ was set to 0, indicating negligible reaction progression and non-specific binding at longer time-points. Second-order association rate constant *k*_on_was derived as:

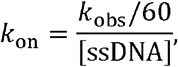

converting *k*_obs_ to *s*^−1^ and dividing by the measured ssDNA molar concentration. *k*_on_ values were adjusted to the actual ssDNA concentrations, which were measured using a NanoDrop spectrophotometer (Thermo Fisher Scientific) by averaging absorbances at 260 nm (for ssDNA) and 647 nm (for 5’-Cy5-label), weighted by ssDNA length. For example: for 14 nt ssDNA (MT002), was weighted as 20% to the 260 nm signal and 80% to the 647 nm signal, and 90 nt ssDNA (MT006) was weighted as 50% to each signal. Experimentally measured correction factors were applied to obtain the final rates for each condition.

### SYTOX Orange intercalation modelling

The approach extends the torsionally constrained intercalator binding model developed by Kolbeck et al.(*27*), which integrates the McGhee-von Hippel (MvH) site-exclusion model with torque-dependent binding energetics for DNA. This framework was extended to back-calculate the initial dye concentration required to achieve a target supercoiling density (σ) after equilibration to a fixed final concentration, accounting for finite-concentration effects and iterative torque equilibration.

MvH model was employed for binding to torsionally relaxed DNA which accounts for cooperative exclusion of binding sites due to the dye footprint. The fractional occupancy γ (dye/bp) is given by:

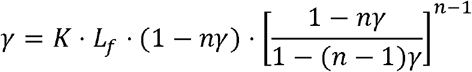

where *K* is the intrinsic association constant (M□^1^), *n* is the binding site size (bp/dye), and *L*_*f*_ is the free dye concentration (M). For finite total DNA and dye concentrations, *L*_*f*_ = *C*_*d*_ − *γ* · *C*_*DNA*_, where *C*_*d*_ is the total dye concentration (M) and *C*_*DNA*_ is the total DNA concentration (molarity of base pairs). This equation was solved numerically for γ using the *fsolve* function from *scipy* optimize module, with γ clamped between 0 and 1/*n* to ensure physical validity.

Parameters for SYTOX Orange were: *K* = 2.4 × 10^6^ M□^1^, *n*= 3 bp/dye, unwinding angle Δ*Tw* = 19.1 degree/dye, elongation Δ*z* = 0.3 nm/dye. DNA length was *N*_*bp*_ = 41,364 bp (corresponding to the uncut plasmid pMT004), with calculated linking number *Lk*_0_= *N*_*bp*_ /10.5 ≈ 3,940 turns. Simulations assumed single-molecule conditions with *C*_*DNA*_ = 10^−12^ M base pairs determined by approximating ~100 double-tethered molecules per field of view (FOV) and scaling up to the total bottom surface of the flow cell for reach *n* (number of molecules), and further calculating the total volume of the flow cell using 0.5 mm as flow cell height (double-sided tape thickness). For torsionally constrained DNA, an iterative solver based on Kolbeck et al(*27*) was implemented. The total linking number difference Δ*Lk* partitions into twist (Δ*Tw*) and writhe (Δ*Wr*) as Δ*Lk* = Δ*Tw* + Δ*Wr* which was calculated using the assumption of 20% of Δ*Lk* residing in twist, thus ((*ω*_*tw*_ = 0.2), with the remainder 80% in writhe.

The equilibrium torque Γ(pN·nm) arises from twist strain:

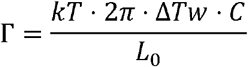

where *kT* = 4.1pN·nm (thermal energy at room temperature), *C* = 100nm (torsional persistence length), and *L*_0_ = 0.34 · *N*_*bp*_ nm (initial contour length). Binding unwinds DNA by Δ*θ* = Δ*Tw* · *π*/180radians/dye and elongates it by Δ*z*/dye, updating *L*_*c*_ = *L*_0_ + *N*_*b*_ · Δ*z*, where *N*_*b*_ is the number of bound dyes.

Torque affects the dissociation constant *K*_*d*_ = 1/Kvia the free energy change per bound dye:

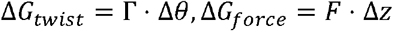

yielding a torque-adjusted 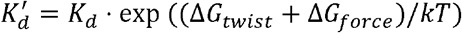, where *F* is applied force (set to 0 pN). Negative torque (underwinding of DNA) favors intercalation by reducing Δ*G* _*twist*_ during additional intercalations. The solver thus initializes with MvH binding at zero torque (*γ*_0_), then iteratively updates *N*_*b*_.

The next aim was to compute the resulting σ of topologically-constrained surface-tethered molecules, which is reached equilibrium after washing SxO from the flow cell. To determine the initial SxO concentration needed to achieve a target σ at a fixed final concentration (Fig. S2B), this was solved inversely. Supercoiling density was calculated as σ = Δ*Lk*/*Lk*_0_, with target Δ*Lk*_*target*_ *SyO* = *σ*_*target*_ *Lk*_0_. For relaxed DNA (σ = 0), [SxO_init_] is equal [SxO_final_]. For negative σ, an initial binding phase at [SxO_init_] induces topological unwinding Δ*Lk*_*topo*_ = − *α* · *N*_*open*_, where *α* = Δ*Tw*/360 and *N*_*open*_ is the pre-bound dye amount from the MvH model at the concentration of [SxO_init_]. Washing the flow cell at [SxO_final_] partially releases dyes, reaching a new equilibrium *N*_*final*_ via the supercoiled model with initial Δ*Lk*_*topo*_. The effective Δ*Lk*_*eff*_ = Δ*Lk*_*topo*_ + *α* · *N*_*final*_, and *σ*_achieved_ = Δ*Lk*_*eff*_ /*Lk*_0_.

The objective *f*([*SxO*]_init_)= Δ*Lk*_*eff*_ − Δ*Lk*_*target*_ was minimized using *fsolve*, with initial guesses: *N*_*open,guess*_ = |Δ*Lk*_*target*_ |/*α, γ*_*guess*_ = *N*_*open,guess*_ /*N*_*bp*_, and free dye approximated inversely from MvH. Unphysical solutions were excluded by bounds (0 mM < [SxO_init_] < 100 mM). Retention (%) = *N*_*final*_ /*N*_*open*_ × 100, with saturation noted if [SxO_init_] > μ100 μM. Simulations scanned [SxO_init_] from 1– 2,000 nM [SxO] which is shown in Fig. S2A-B. Our model here assumed uniform torque, no buckling transitions, and negligible force (F = 0), focusing on ensemble-averaged behavior under single-molecule conditions. Further assumptions might be necessary when applying this model to more extreme σ values where e.g., negatively supercoiled DNA molecules can experience local denaturation and strand separation.

### RecA filament sampling reach and radius-of-gyration (R_g_) estimate

We estimated the length of dsDNA sampled by a RecA presynaptic filament by first computing its contour length as *L*_fil_ = *N*_nt_ × 0.51 nm, where the axial rise of 0.51 nm per nucleotide is the established value for the ATPγS-stabilized RecA–ssDNA filament(*38*). Radius of gyration was calculated by treating the rigid filament as a straight rod, *R* _*g*_ = *L*_fil_ /2. Here, *R*_*g*_ was converted into an equivalent random-coil dsDNA contour length using the WLC-like Gaussian chain approximation valid for 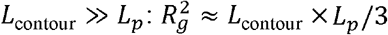 with dsDNA persistence length *L*_*p*_ = 50 nm. The corresponding number of base pairs sampled was converted back from physical distance as *N*_*bp*_ = *L*_contour_/0.34. We performed these calculations for invasion strand lengths from 1 to 600 nt shown in Fig. S4C.

### Statistical analysis

No statistical method was used to predetermine sample size. For single-molecule experiments, individual DNA molecules were treated as individual observations, and experiments were repeated across independent flow cells and imaging days. For live-cell experiments, individual cells were treated as individual observations, and experiments were repeated across independent cultures. The exact number of molecules, cells, independent experiments and biological replicates is reported in the corresponding figure legends.

Single-molecule binding curves were generated from the averages of a minimum of three fields of view over two independent flow cells (experimental repeats). The curves were fitted to the average values of bound fraction (individual cy5-ssDNA-RecA filament to dsDNA on the surface). Apparent association rates were obtained by fitting binding curves to a single-exponential association model, unless stated otherwise. Fitted parameters are reported as mean ± SD or mean ± 95% confidence interval, as indicated in the figure legends. DNA end-to-end distances, RecA filament positions, plectoneme positions and live-cell RecA-YFP trajectories were extracted using custom python image-analysis as described above.

### Caulobacter crescentus strains and growth conditions

Strains and plasmids used in this study are listed in Tables S4-S5, respectively. Chromosomal engineering (such as insertion of fluorescent markers) was carried out either by the standard two-step recombination approach or using previously described vectors(*70*) and details of the same are provided in the respective tables. Transductions were performed with ΦCr30(*71*). *Caulobacter* cultures were grown at 30°C in peptone yeast extract (PYE) medium and supplemented with antibiotics at the appropriate concentrations.

For strains in which *dnaA* was placed under an IPTG-inducible promoter, liquid cultures were supplemented with 0.5mM IPTG and agar plates with 1mM IPTG. For synchronization experiments, we grew bacterial cultures to mid-log phase and synchronized as previously described(*11, 45-47*). Briefly, to isolate non-replicating pre-divisional cells, bacterial cultures were grown to mid-log phase in IPTG-containing medium, synchronized, and swarmer cells were isolated. The swarmer cells were released into medium lacking IPTG, depleted of DnaA by washing out the inducer, and grown for one generation in the absence of IPTG. Cell division was then halted by addition of 35 μg/ml cephalexin, and the resulting non-replicating pre-divisional cells were used for DSB induction experiments.

In all cases, recA-YFP was induced with 0.03% xylose 90 min prior to imaging, and ParB-YFP was induced with 0.3% xylose 3 h prior to imaging. DSBs were induced with 2 µM vanillate 15 min prior to imaging, and the same concentration was maintained in the agarose pad throughout imaging. For experiments with novobiocin, cells were exposed to 50 µg/ml novobiocin for 30 min at the time of DSB induction, prior to time-lapse imaging. For recovery experiments, novobiocin-treated cultures were washed twice in fresh PYE and released into fresh medium containing vanillate and xylose to induce I-SceI and *recA* expression, respectively. Cells were imaged shortly after release into fresh medium. All inducers were maintained at the same concentrations in the agarose pads during imaging.

### Fluorescence microscopy and image analysis

For snapshot microscopy, aliquots were withdrawn from cultures grown under the indicated conditions at defined time points, pelleted, and resuspended in an appropriate volume of growth medium. A 2 µl drop of the resulting cell suspension was applied to 1% agarose pads (Invitrogen ultrapure; ~0.5 × 0.5 cm). For time-lapse imaging, cells were imaged on 1.5% GTG (low-melting) agarose pads prepared in peptone yeast extract and supplemented with vanillate, xylose, cephalexin or novobiocin at the appropriate concentrations, using glass-bottom Petri dishes. Imaging was performed on a Nikon Eclipse Ti2 wide-field epifluorescence microscope equipped with a motorized XY stage, using a 60x Plan Apochromat objective (NA 1.4) and illumination from a pE4000 light source (CoolLED). Images were acquired with a Hamamatsu ORCA-Flash 4 camera using Nikon NIS-Elements software. Focus was maintained throughout time-lapse acquisition using the infrared-based Perfect Focusing System. For fluorescence microscopy, GFP, YFP, and mCherry were excited at 470 nm, 490 nm, and 550 nm, respectively. For all experiments with RecA, exposure time used for excitation at 490 nm was 400 ms. For imaging HupA, exposure time used for excitation at 550 nm was 100 ms. For imaging ParB, exposure time used for excitation at 490 nm was 400 ms. Images were collected every 10 sec, 2 min, or 5 min (as specified in the main text and corresponding figure legends). Images were analyzed using MicrobeJ(*72*), Oufti, or by manual assessment in ImageJ as previously described(*11*). All experiments were performed with 2-3 biological replicates, and data from at least 2 independent repeats were pooled and plotted after confirming reproducibility across replicates. Plots were generated and statistical analyses were carried out using R or GraphPad Prism. Schematics were prepared in Adobe Illustrator.

## Acknowledgments

We thank Dr. Brandon Case, Dr. Joshua Cofsky, Gemechu Mekonnen, Dr. Jakub Wiktor and Dr. Jovana Kaljević for discussions and advice for this work. We also thank Dr. Afroze Chimthanawala for initial assistance with reagent generation.

## Funding

This publication and project results were supported by Dutch Research Council (NWO) under the NWO Rubicon program awarded to M.T (project number 019.242EN.011). S. B was supported by a graduate student fellowship from the Tata Institute of Fundamental Research. The project was also supported by a grant from the National Institutes of Health (R35GM158214) to J.J.L and by the Department of Atomic Energy, Government of India, Project Identification number RTI 4006 to A. B.

## Author contributions

Conceptualization: M.T., A.B., J.J.L.

Methodology: M.T, S.B, S.C.

Experimental investigation: M.T, S.B.

Formal analysis: M.T, S.B.

Visualization: M.T, S.B.

Funding acquisition: M.T., T.H, A.B., J.J.L.

Supervision: T.H, A.B., J.J.L.

Writing - Original Draft: M.T, A.B.

Writing - review & editing: M.T, S.B, S.C, T.H, A.B, J.J.L.

## Data availability

All data mentioned in this work is shown in the Figures. All raw data from experiments are available upon request to corresponding authors. Correspondence and requests for materials should be addressed to A.B. and J.J.L.

## Code availability

Analytical model for supercoiling density calculation is available at the GitHub repository: [https://github.com/milostisma/Supercoiling-calibration-model].

## Competing interests

The authors declare no competing interests.

## Notes

### Competing Interest Statement

The authors have declared no competing interest.

