## Supplementary figures for "Chromosome topology gates productive RecA homology search"

**This supplementary section includes:**

Supplementary Figures S1 to S8

Supplementary figure legends

Supplementary Movies S1 to S5 legends

Supplementary Tables S1 to S5

### Supplementary Figures

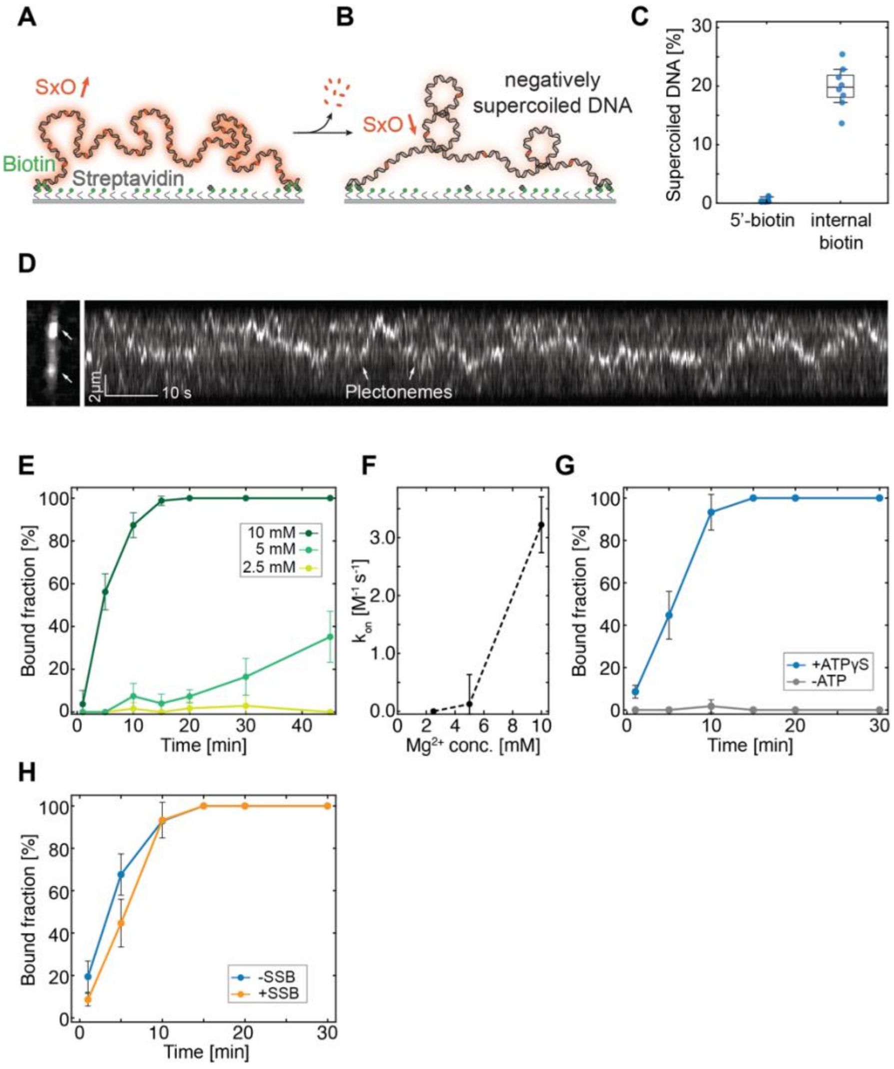

**Figure S1. Construction of supercoiled, surface-tethered DNA construct for homology searc**

**A)** Schematic representation of single-molecule surface assay. Long 42kb DNA is attached to the surface at high SYTOX Orange concentrations, via multiple-biotin streptavidin interactions to ensure topological constraint in an underwound state. **B)** Topologically constrained DNA is washed of excess SxO intercalating dye, which results in negatively supercoiled DNA. **C)** Direct comparison between fractions of supercoiled DNA molecules in the presence of a single 5'-biotin and multiple internal biotins incorporated using a PCR in the presence of Biotin-dUTP and dTTP mixture (see Methods). Each datapoint represents an average value from an individual FOV (n > 20 for each). **D)** Left – a snapshot of a supercoiled, surface-tethered DNA molecule stained with SxO. Right - Kymograph of the same molecule with diffusing plectonemes. White arrows show DNA plectonemes, appearing as DNA puncta of higher intensity. **E)** Timecourse experiments during the flow cell incubation with 10nM ssDNA-RecA filament, in the varying $Mg^{2+}$ concentrations (10mM: n = 479, 5mM: n = 439, 2.5mM: n =

450). Error bars represent standard deviation for each time point measurement. **F)** Binding rates ($k_{on}$) calculated by fitting the curves show in E) with a pseudo-first order kinetic binding model. Error bars represent the standard error of the fitted pseudo-first-order rate constant. **G)** Timecourse experiments during the flow cell incubation with 10nM ssDNA-RecA filament, in the presence or absence of 1 mM ATP/ATPγS (+ATPγS: n = 485, -ATP: n = 327). **H)** Timecourse experiments during the flow cell incubation with 10nM ssDNA-RecA filament, in the presence of 20nM SSB protein (n = 308). Error bars represent standard deviation for each time point measurement.

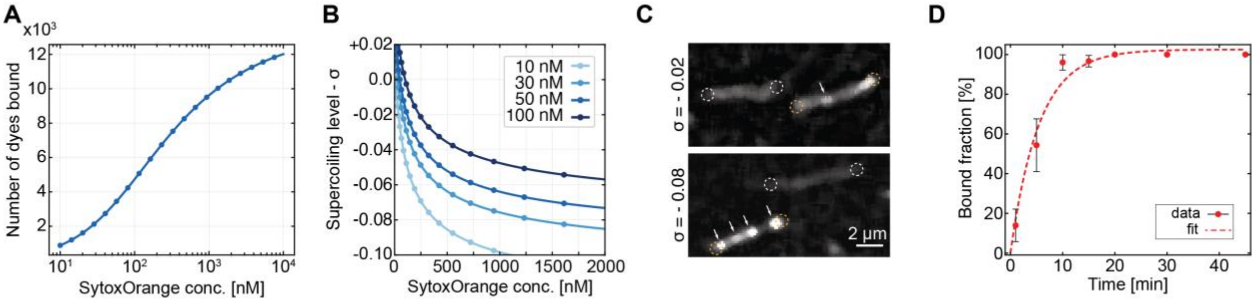

**Figure S2. Analytical model of SxO intercalation and supercoiling density calculations**

**A)** Non-linear dependence of SxO intercalation at different SxO concentrations. For reference: we used the regime of 260 nM to 50 nM, in most experiments where σ = -0.05. **B)** Analytical calculation of starting SxO concentrations (before torsionally constraining) required to induce a specific supercoiling density after washing at either 10, 30, 50 or 100nM SxO concentration. **C)** Micrographs showing DNA molecules at different supercoiling density (Top – at σ = -0.02, and bottom – at σ = -0.08) in comparison to non-coiled DNA molecules. White arrows indicate plectonemes. **D)** Pseudo-first order reaction fit to the timecourse data for σ = -0.08 shown in Fig. 1E, which was used to calculate the $k_{on}$ rates in Fig. 1F. Error bars represent standard deviation for each time point measurement.

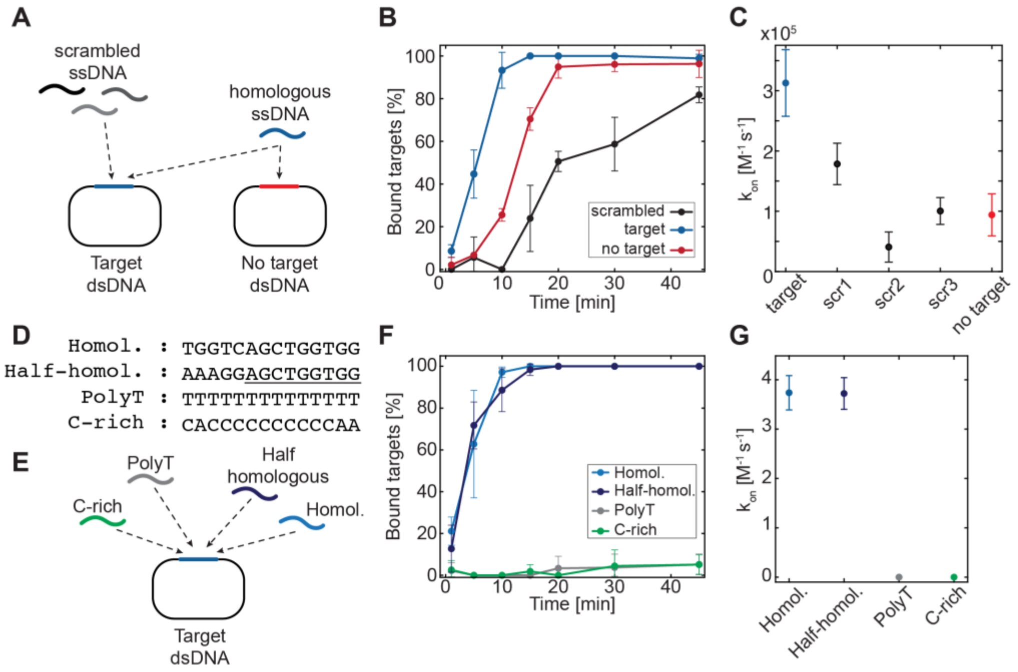

**Figure S3. Full homology to target molecules ensures highest binding rate during homology search**

**A)** Schematic representation of the timecourse experiment in panel B, where heterologous binding was measured using 90-nt invasion strands with ~50% GC content that do not share extended homology with the surface-tethered dsDNA. Presynaptic filament with the same ssDNA (90nt homologous – in blue), was tested against surface tethered dsDNA lacking the 90 bp homologous region (in red) while keeping the entirety of 42 kb sequence identical. **B)** Timecourse experiments using non-homologous invasion strands at 10 nM ssDNA-RecA filament. $n_{(target)}$ = 642 (same data as Fig. 1D), $n_{(scrambled)}$ = 526, $n_{(no\ target)}$ = 375. **C)** Binding rates ($k_{on}$) calculated by fitting the curves shown in B) with a pseudo-first order kinetic binding model. Error bars represent the standard error of the fitted pseudo-first-order rate constant. **D)** ssDNA constructs used in 14nt minimal filaments to compare absolute requirements for microhomology and possible non-specific binding. **E)** Schematic representation of the target molecule (in a plasmid, rather than surface tethered form) and different minimal 14nt RecA filaments used in the timecourse experiments. **F)** Timecourse experiments during the flow cell incubation with 10nM ssDNA-RecA filament, with the varying ssDNA invasion strands (Homologous: n = 532 (same as in Fig. 1G), partially homologous: n = 487, T-rich: n = 282, C-rich: n = 355). Error bars represent standard deviation for each time point measurement. **G)** Binding rates ($k_{on}$) calculated by fitting the curves show in F) with a pseudo-first order kinetic binding model. Error bars represent the standard error of the fitted pseudo-first-order rate constant. Binding rates for curves that do not reach 10% total target binding over 45 min, are pinned to 0 $M^{-1}s^{-1}$.

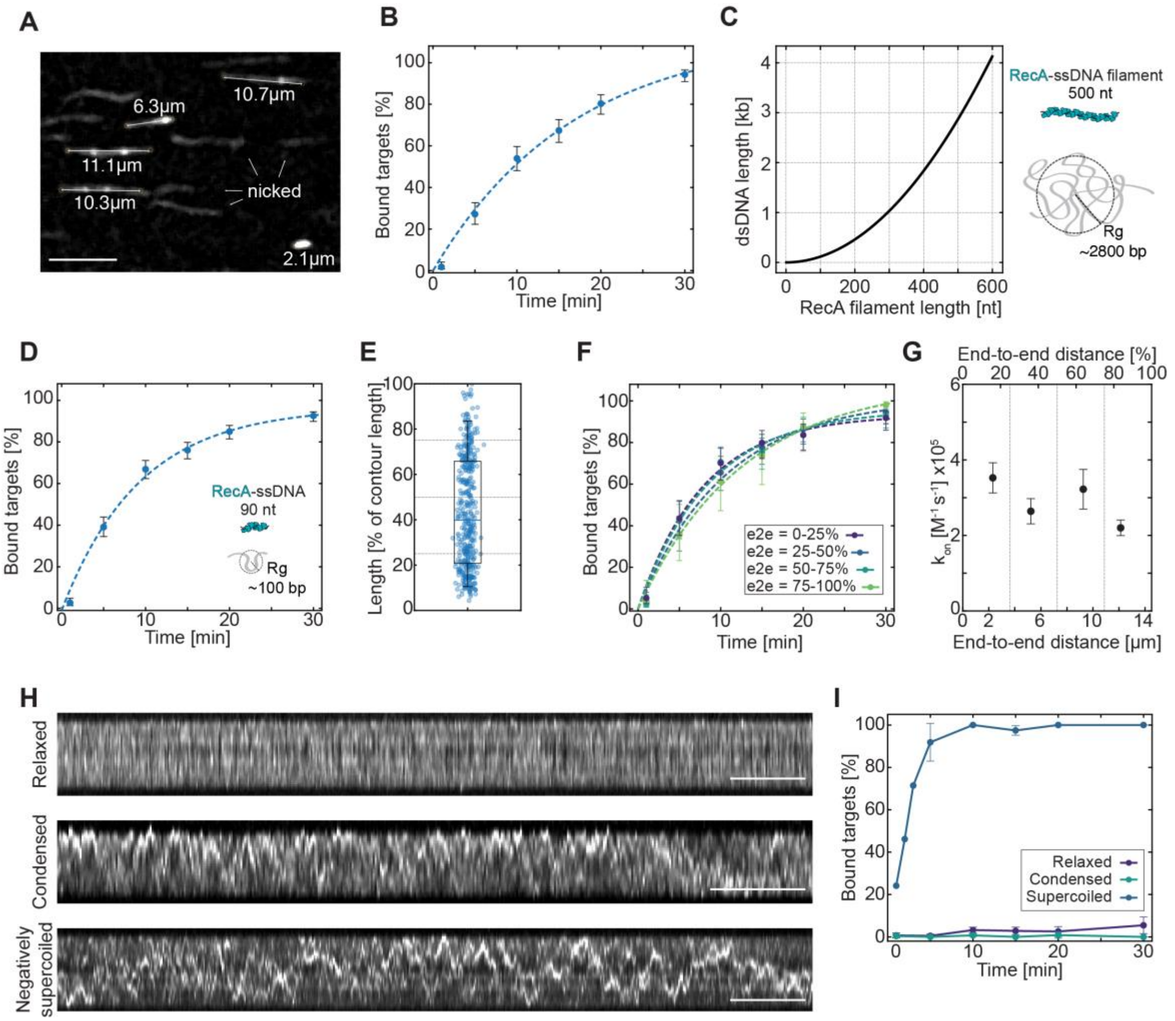

**Figure S4. DNA compaction of surface-bound molecules does not affect homology binding**

**A)** Micrographs showing DNA molecules at different end-to-end tethering distance within the same flow cell and field of view. White arrows point to nicked, non-coiled DNA molecules**.** Scale bar – 10 µm **B)** Aggregate timecourse binding curve of all molecules shown in Fig. 2B, C. Blue dashed line represents a pseudo-first order reaction fit to the data. Error bars represent a 5-95$^{th}$ percentile confidence interval for a binomial distribution at each timepoint. Total n = 279 for each time point. **C)** Predicted reach of RecA filaments relative to untethered double-stranded DNA. Radius of gyration ($R_g$) is calculated based on WLC-like gaussian-chain model of the DNA (see *Methods*). RecA filament length is calculated as an average of 0.51 nm/nt in a fully assembled filament(*1*). **D) -G)** Same as Fig. S4B, Fig. 2B, C, D, respectively but for 90 nt RecA filament. Total molecules n = 434 ($n_{(0\text{-}25\%)}$ = 134, $n_{(25\text{-}50\%)}$ = 125, $n_{(50\text{-}75\%)}$ = 49, $n_{(75\text{-}100\%)}$ = 126). **H)** Kymographs of relaxed surface-tethered DNA molecules (top), condensed DNA molecule (7.5 % PEG 8000, middle), and negatively supercoiled DNA molecule (σ = -0.05) shown in Fig. 2E. Scale bars = 10 s. **I)** Timecourse experiments using 500 nt RecA filament at 0.2 nM concentration. $n_{(relaxed)}$ = 1281, $n_{(condensed)}$ = 1579, $n_{(supercoiled)}$ = 603 (same data as Fig. 1G - 500 nt).

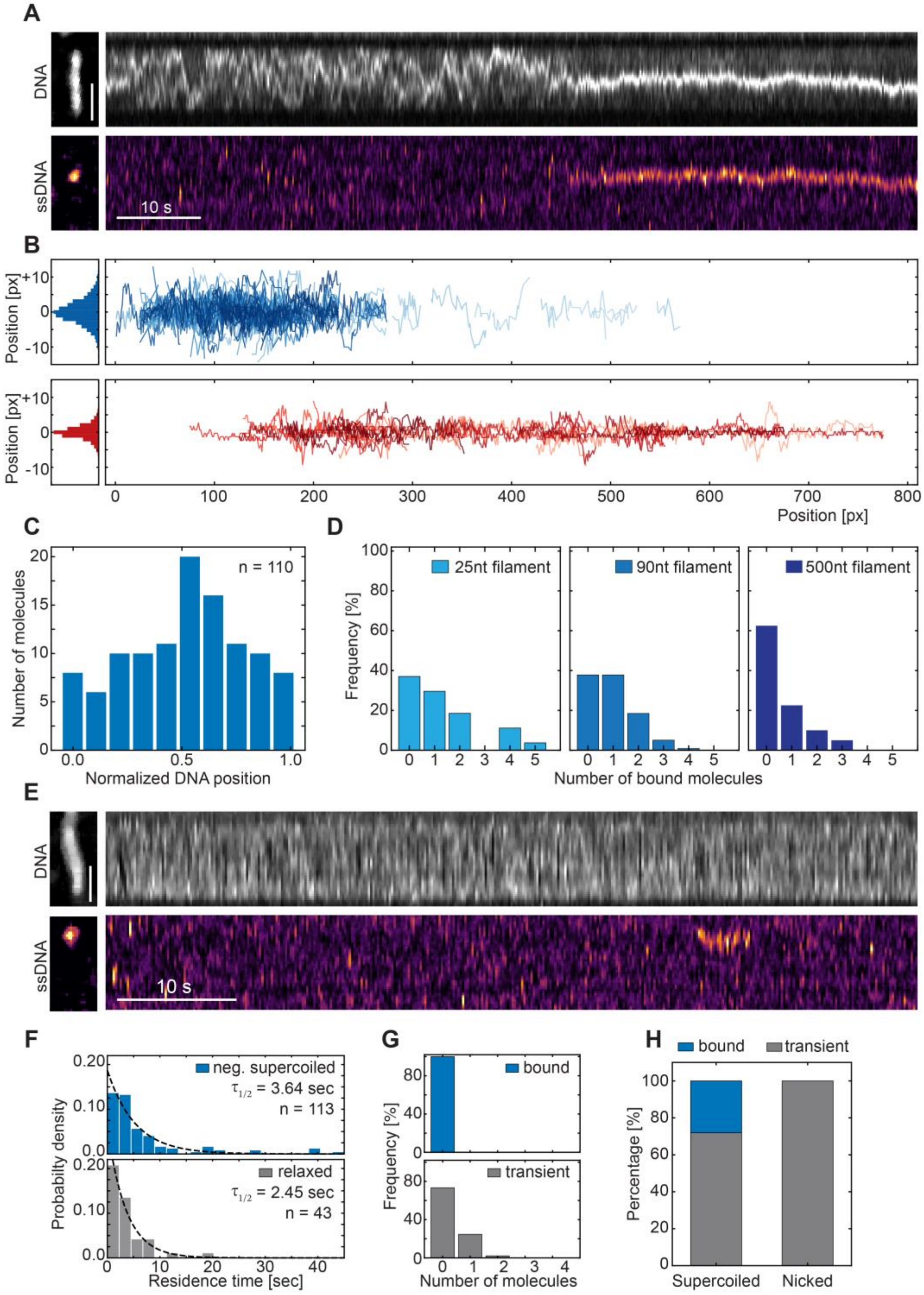

**Figure S5. Transient sampling and stable binding define two modes of RecA filament interaction with DNA**

**A)** Top – Kymograph SxO-stained surface tethered dsDNA with a corresponding DNA molecule micrograph on the left. Bottom – kymograph for Cy5-labelled ssDNA-RecA presynaptic filament with a corresponding micrograph of the same region as above, in the moment of DNA binding. Scale bar – 2µm. **B)** DNA plectoneme dynamics tracked by the peak position movement over time. Top/Blue – overlay of 207 DNA plectoneme tracks prior to RecA-ssDNA filament binding. Corresponding histogram showing the lateral coverage of the plectoneme diffusions (centered to zero at average positions) shown on the left. Bottom/Red – overlay of 46 pinned plectoneme tracks after RecA-ssDNA filament binding with a corresponding histogram on the left. **C)** Frequency of transiently bound molecules to dsDNA targets on the surface relative to RecA filament length. ($n_{(25\ nt)}$ = 46, $n_{(90\ nt)}$ = 157, $n_{(500\ nt)}$ = 46). **D)** Off-target binding positions of transiently bound 90 nt RecA filaments to supercoiled surface-tethered DNA molecules (n = 110 binding events). See Fig. 3G for transient binding on supercoiled molecules. **E)** Same as Fig. 3G, but for a relaxed DNA molecule. Scale bar – 2µm. **F)** Residence time histograms of transiently bound RecA filaments to supercoiled DNA targets (top/blue, n = 113, same as Fig. 3H) and relaxed DNA targets in the same experiment (bottom/gray, n = 43). Black dashed lines represent exponential decay fit to histogram data. **G)** Frequency of transient (gray/bottom) and stable (blue/top) binding to relaxed surface-tethered DNA target molecules. Same as Fig. 3F but for relaxed DNA molecules. **H)** Direct comparison between the percentage of stably bound (blue) and transiently bound (gray) RecA filaments to either supercoiled (n = 157) or nicked (n = 43) surface-tethered DNA target molecules.

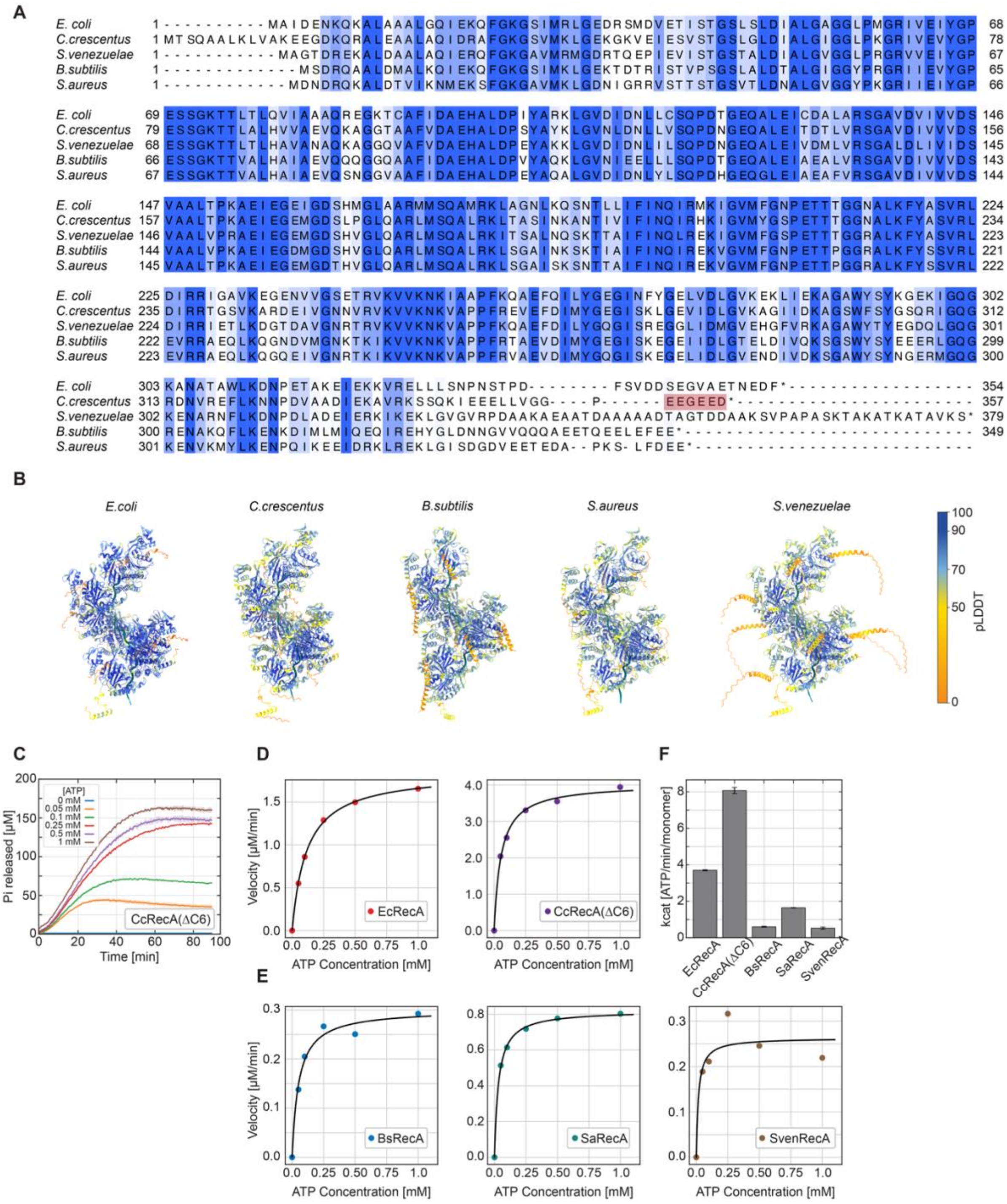

**Figure S6. Structural and biochemical comparison of bacterial RecA homologs**

**A)** Multiple sequence alignment of five RecA homologs (*Escherichia coli, Caulobacter crescentus, Streptomyces venezuelae, Bacillus subtilis, Staphylococcus aureus*). Sequence of RecA protein from *E. coli* was used as reference. Red shaded amino-acid residues in *C. crescentus* RecA (CcRecA), were removed for our single-molecule experiment, as full CcRecA variant did not show any double-stranded DNA binding. **B)** AlphaFold model of 9-mer RecA filament bound to 24 nt ssDNA for each RecA homolog showing right-handed architecture of the RecA filament. **C)** Determination of ATPase activity at different ATP concentrations. Exemplary timecourse measurement of *C. crescentus* RecA mutant (CcRecA(ΔC6)). Light shaded areas of the corresponding colors represent standard deviation from duplicate experiments. **D-E)** ATP hydrolysis velocity obtained as linear fits to data presented in panel C for each RecA homolog. Black dashed lines represent Michaelis-Menten fit to data points. Panel D shows RecA homologs from Gram-negative bacteria, panel E shows RecA homologs from Gram-positive bacteria. All experiments were performed in duplicates (n = 2). Colors correspond to Fig. 4. **F)** Catalytic rates of each RecA homolog obtained from MM fits (in panel D), and adjusted for experimental concentration of 0.5 μM RecA.

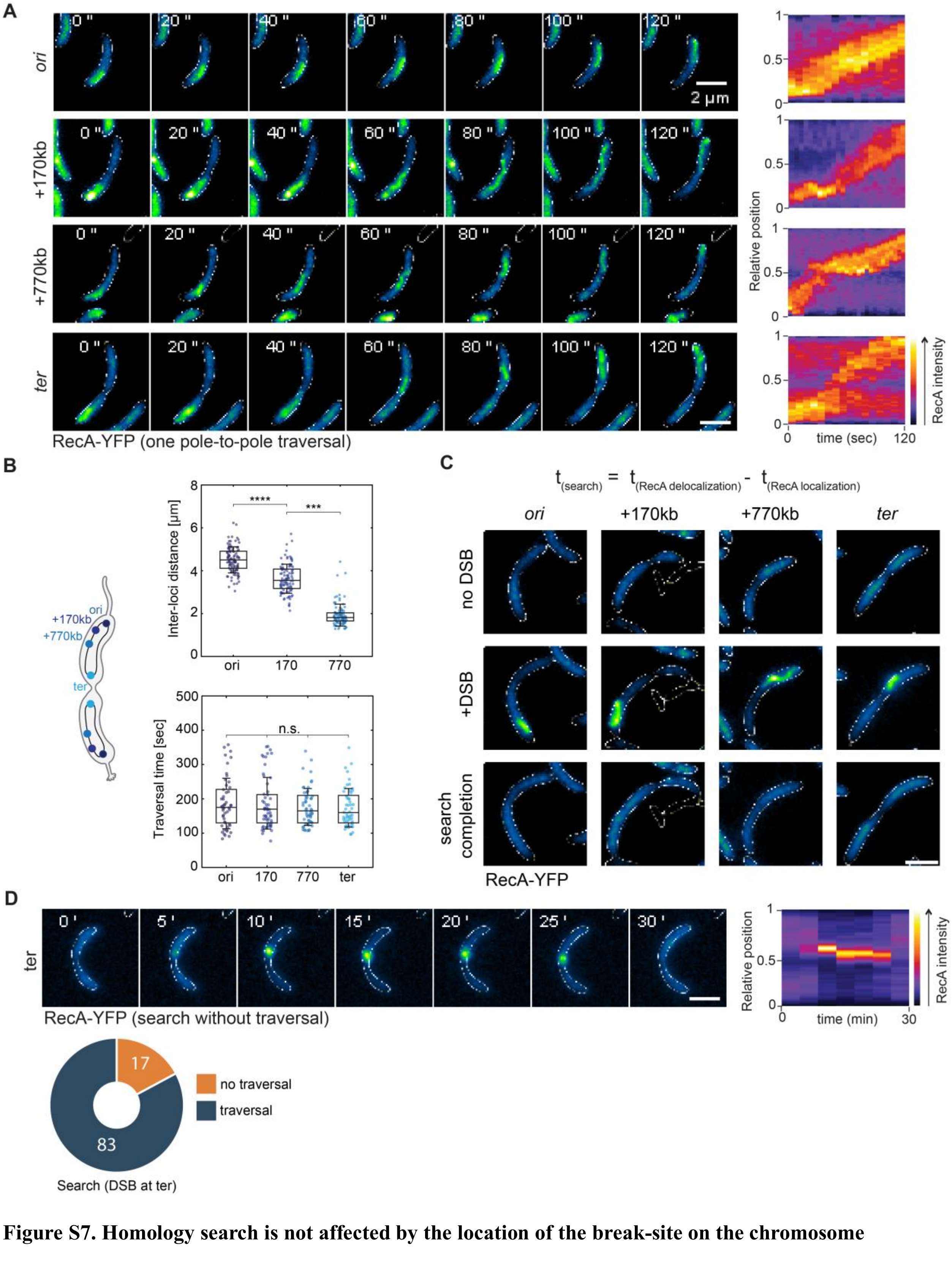

**Figure S7. Homology search is not affected by the location of the break-site on the chromosome**

**A)** Left - Time lapse tracking of RecA filament movement during homology search for DSB induced at different locations (30 kb/*ori*, 170 kb, 770 kb, 1958 kb/*ter*). Scale bar = 2 µm. Times, in seconds, indicated at each micrograph. Right - Kymograph of RecA position normalized to the cell length for the respective montages shown on the left (images taken every 10sec). Same as shown in Fig. 5B. **B)** Top - Inter locus distances between two homologous regions are measured as a proxy for the distance between break site and repair template at different DSB locations. Box plots represent the median, interquartile ranges and the whiskers indicate 10-90 percentile range. Individual dots represent inter locus distances for a single cell. $n_{ori}$ = 100, $n_{170kb}$ = 100, $n_{770kb}$ = 100, pooled from at least 2 independent experiments. Significance score: P < 0.0001, using two sided Mann-Whitney test. Bottom - RecA filament traversal time in cells following DSB induction at different locations (as indicated in panel A). Box plots represent the median, interquartile ranges and the whiskers indicate 10-90 percentile range. Individual dots represent traversal times for a single cell. $n_{ori}$ = 50, $n_{170kb}$ = 56, $n_{770kb}$ = 56, $n_{ter}$ = 53 cells, pooled from at least 2 independent experiments. Significance score: ns, using two sided Mann-Whitney test. **C)** Representative cells with DSB at different locations as indicated in panel A**;** RecA-YFP localization in the absence of DSB, upon DSB induction, and at the completion of search is shown. Search time ($t_{search}$) is calculated as the difference between time at RecA delocalization ($t_{RecA\ delocalization}$) and the time at which RecA localization first appeared ($t_{RecA\ localization}$). Scale bar = 2 µm. **D)** Example a rare case where search is completed without significant long-distance traversal by the RecA filament following DSB induction at *ter.* See Fig. 5C (*ter* - gray labelled data points). Left - Time lapse tracking of RecA filament position in case of homology resolution without traversal. Time in minutes, indicated at each micrograph. Scale bar = 2 µm. Right - kymograph of the RecA-YFP position in the same cell. Pie chart below indicates the percentage of cells where search completion is observed at *ter* either with or without RecA traversals. Mean is shown within each sector of the pie graph. n = 100 cells pooled from at least 3 independent experiments.

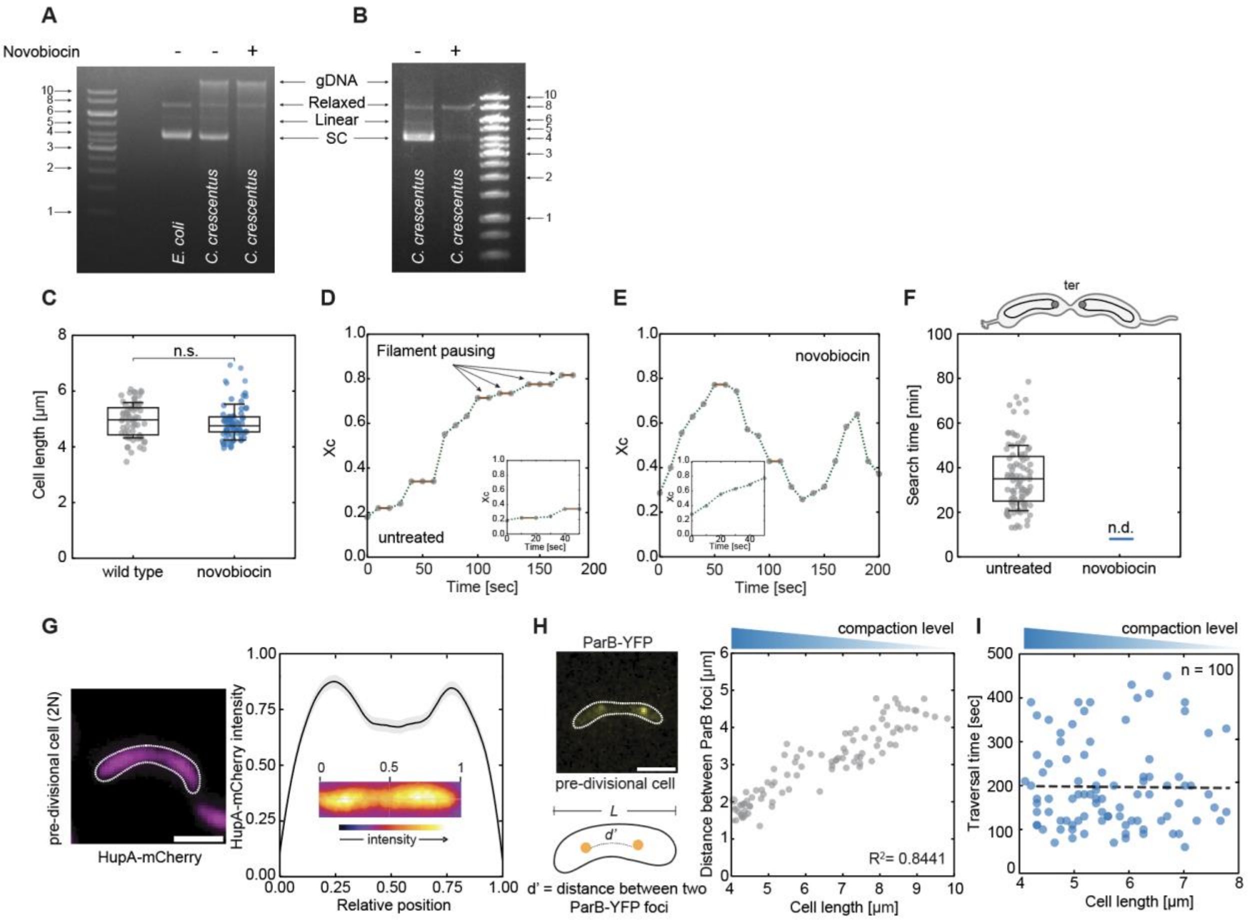

**Figure S8. DNA supercoiling ensures productive homology search *in vivo*.**

**A)** Plasmid species isolated from *C. crescentus* before and after novobiocin treatment (50 µg/ml, 30 minutes). The same plasmid isolated from untreated *E. coli* DH5α was included as a control. Agarose gel electrophoresis was performed followed by ethidium bromide staining. DNA ladder sizes indicated in kb. **B)** Same as panel A, stained with SYBR Gold. **C)** Cell length distributions of untreated (gray, n = 66) and novobiocin-treated (blue, n = 73) cells used in analysis (Fig. 5E-H). Box plots represent the median, interquartile ranges and the 10-90 percentile range. Individual dots represent cell lengths of single cells, pooled from 3 independent experiments. Significance score: ns, using unpaired t test. **D)** Representative tracking example of RecA filament movement in untreated cells. Position calculated as displacement of the filament centroid position - $x_C$ (relative to cell length) measured across time over a single traversal (images taken every 10 sec). Dashed lines indicate periods of directional movement, whereas solid lines indicate pause events. Inset shows the filament movement during a single traversal in novobiocin-treated cells illustrating that untreated cells show longer pause durations. **E)** Representative example RecA filament movement in novobiocin-treated cells (images taken every 10 sec). Multiple traversals are shown over the same time period, due to faster filament movement. Inset shows the centroid position for the time duration of a single traversal in novobiocin-treated cells illustrating that cells rarely show pausing during directional movement. Dashed lines indicate periods of displacement, whereas solid lines indicate rare pause events. **F)** Total homology search time of RecA filaments in untreated (gray, n = 100) and novobiocin-treated cells (blue/n.d., n = 0/100) when DSB is induced at *ter* (see Fig. 5H for comparison). Box plots represent the median, interquartile ranges and the 10-90 percentile range. Individual dots represent search times of single cells pooled from at least 2 independent experiments. Data for homology search time of RecA filament in untreated cells at ter has been reproduced from Fig. 5C for comparison. **G)** Left - localization of HupA-mCherry (purple) in a representative pre-divisional cell. Scale bar = 2 µm Right - Intensity profile of HupA-mCherry across the cell length. Inset shows intensity heatmap of the same cell. **H)** Inter-locus distance (d') between two homologous loci (+770kb) in pre-divisional cells is calculated. The loci are marked by a *parS-pMT1* site bound by ParB-pMT1-YFP. Inter-locus distance is plotted as a function of cell length. n = 97 (from 2 independent experiments). Pearson's correlation coefficients ($R^2 = 0.8441$) with CI (95% [0.881-0.945]). Observed degree of change in chromosome compaction is represented as a gradient blue triangle (top). **I)** RecA filament traversal time relative to *C. crescentus* cell length. Linear fit correlation: $R^2 = 0.0003$, n = 100. Black dashed line represents a linear regression fit to data.

**Supplementary Movie 1. Dynamics of negatively supercoiled DNA molecules in the single-molecule assay.** Supercoiling density (σ) of -0.02, -0.05, and -0.08 shown. White arrows indicate negatively supercoiled molecules in each panel. Scale bar = 2 µm.

**Supplementary Movie 2. RecA-YFP filament homology search in untreated *C. crescentus* cells.** Time, in minutes, indicated on the top. Scale bar = 2 µm

**Supplementary Movie 3. RecA-YFP filament homology search in untreated *C. crescentus* cells.** Time, in minutes, indicated on the top. Scale bar = 2 µm

**Supplementary Movie 4. RecA-YFP filament homology search after novobiocin treatment.** Time, in minutes, indicated on the top. Scale bar = 2 µm

**Supplementary Movie 5. RecA-YFP filament homology search after novobiocin treatment.** Time, in minutes, indicated on the top. Scale bar = 2 µm

| **Primer name** | **Sequence** |
|---|---|
| SC2080 | AAATAATTTTGTTTAACTTTAAGAAGGAGATATACatgGCTATCGACGAAAACAA |
| SC2081 | TCCTGGCTGTGGTGATGATGGTGATGGCTGCTGCCttaAAAATCTTCGTTAGTTT |
| T-21 | ATACCTACAGCGTGAGCTATGAGAAAGC |
| T-22 | TTCCGCCGTATACGTTTGCCAGC |
| T-43 | **5'-Bio-**ATACCTACAGCGTGAGCTATGAGAAAGC |
| T-44 | **5'-Bio-**TTCCGCCGTATACGTTTGCCAGC |
| T-46 | **5’-Phos**-TAGGGCAGCAGCCATCACCA |
| T-49 | **5’-Phos**-AGGGCCTCCTACCAACAGCTC |
| DR_16 | CAAGCTTCTCTGCAGGATATCTGGACGACGACGCCCCCTGAGG |
| DR_17 | CCGGAGCTCGAGATCTTAAGGTACCGCCGTTGACGGCGTCCTTCAGTTG |
| DR_09 | GGTACCTTAAGATCTCGAGCTCCGGAGAATTCG |
| DR_10 | TTACTTGTACAGCTCGTCCATGCCGC |
| DR_18 | GGCGGCATGGACGAGCTGTACAAGTAAGCCTTCAGAGAGGGGCCGG |
| DR_19 | CGGAGACGCGTCACGGCCGAAGAGCTCGATATTGATCGACACCCGATCG |
| AC_180 | TTAACATATGATGACAAGTCAGGCGGCTTTGAAACTCGTGGCCA |
| AC_202 | TTAAGGTACCGTCCTCTTCGCCCTCTTCCGGGCCGC |
| AB_239 | TTATGGATCCCCAGTTCTTCTGGGCGGCCAC |
| AB_248 | GAACCCGACCGATAGGCCCGCTCAGACGAGGGTTTCGAGCGC |
| AB_184 | CGGTATCGATAAGCTTGATATCGAATTCATGAAG |
| AB_185 | CTGCAGGAATTCAAGAAGTTCCTATTCTCTAGAAAG |
| AB_242 | GGCAACGAGCCGATCGCTGATTCCATCACCCATGTGATTAGCGCC |
| AB_243 | TTATGCTAGCCGCAGCTCGATCGCCATGG |

**Table S1. DNA oligonucleotides used in the presented study for cloning and plasmid construction**

| **Name** | **Sequence** | **Label** |
|---|---|---|
| MT-001 | GCTGGTG | **5'-Cy5** |
| MT-002 | TGGTCAGCTGGTGG | **5'-Cy5** |
| MT-003 | TATTAATGCATATATAGTATCGCCGAACGATTAGCTCTTCAGGCTTCTG AAGAAGCGTTTCAAGTACTAATAAGCCGATAGATAGCCACGGACTTC GTAGCCATTTTTCATAAGTGTTAACTTCCGCTCCTCGCTCATAACAGA CATTCACTACAGTTATGGCGGAAAGGTATGCATGCTGGGTGTGGGGA AGTCGTGAAAGAAAAGAAGTCAGCTGCGTCGTTTGACATCACTGCT ATCTTCTTACTGGTTATGCAGGTCGTAGTGGGTGGCACACAAAGCTT TGCACTGGATTGCGAGGCTTTGTGCTTCTCTGGAGTGCGACAGGTTT GATGACAAAAAATTAGCGCAAGAAGACAAAAATCACCTTGCGCTAA TGCTCTGTTACAGGTCACTAATACCATCTAAGTAGTTGATTCATAGTG ACTGCAGTCTGATCCTTTGCGAATACGCTGGTGGTCAGCTGGTGGGC CCACGCGATGGGTAACAGTCGCTGGTGG | / |
| MT-004 | TCAGCTGGTG | **5'-Cy5** |
| MT-005 | ACCTATCGGCACAAAAGTGACACGACGTCGATGAGTAGCGGACTTTT GGTCAACCACAATTCCCTAAGGGACAGGTCCTGCGGTGTACAT | **5'-Cy5** |
| MT-006 | AGTTGATTCATAGTGACTGCAGTCTGATCCTTTGCGAATACGCTGGTG GTCAGCTGGTGGGCCCACGCGATGGGTAACAGTCGCTGGTGG | **5'-Cy5** |
| MT-010 | AAAGGAGCTGGTGG | **5'-Cy5** |
| MT-011 | GTAAGTGATCTAACGCTTCGGATATGACTATATACTTAAGCTTGATCTC GTCCCGAGAATTCTAAACCTCAACATTTATAGATTATAAGG | **5'-Cy5** |

| | | |
|---|---|---|
| MT-015 | ACCAGTTAACCTCGCAAGAACGTCCTTAGCTCTTGCAGGCAATTAAG GAGAACGTAAGTATAGCGCAATTTAACAGAGAATTAGGTTGAT | **5'-Cy5** |
| MT-016 | CACCCCCCCCCCAA | **5'-Cy5** |
| MT-017 | TTTTTTTTTTTTTT | **5'-Cy5** |
| MT-018 | CTGCAGTCTGATCCTTTGCGAATAC | **5'-Cy5** |

**Table S2. ssDNA used in the presented study presynaptic filament construction**

| **Name** | **Sequence** |
|---|---|
| 540 bp insert | tctgatcctttgcgaatacgcTGGTGGTCAGCTGGTGGgcccacgcgatgggtaacagtcGCTGG TGGttcgcGCGCGAAATTATGAGTCACGAAGAGGTTGAAAAGCGTGGTGA TTTCAGTGTGAAAGAATTTTCAATTTTCCTTTATAATCAAACAAATAACC ATGAAATTGGCGTGGTGAAAAACATACAAAAAAGATGCTCTTCGGCATC CTGAATTCCCGACAGTAAGACGGGTAAGCCTGTTGATGATACCGCTGCC TTACTGGGTGCATTAGCCAGTCTGAATGACCTGTCACGGGATAATCCGA AGTGGTCAGACTGGAAAATCAGAGGGCAGGAACTGCTGAACAGCAAA AAGTCAGATAGCACCACATAGCAGACCAGATCTttaacccaccccgggttacgtaaata ctggcaggcgtttcgtAATTGTGAGCGCTCACAATTgactgccgtccggcctaagcagAGATC CGACGACTCCTAGTTACGCTAGGGATAACAGGGTaattgtgagcggataacaatttg |

**Table S3. DNA sequence used in the presented study for plasmid construction.** Underlined sequence represents homology to the NdeI-linearized plasmid.

| **Plasmid** | **Construction details / Source** | **Antibiotic** |
|---|---|---|
| pMT004 | pBS-chi3 - Described in details above. | Amp |
| pMT007 | HiFi-reaction between NcoI-cut pETDuet1 and SC2080/2081 amplicon from *E. coli* MG1655 genome. | Amp |
| pMT011 | HiFi-reaction between NcoI-cut pETDuet1 and geneblock carrying RecA (*C. crescentus*) sequence Uniprot / B8H3I7 | Amp |
| pMT012 | HiFi-reaction between NcoI-cut pETDuet1 and geneblock carrying RecA (*B. subtilis*) sequence / Uniprot P16971 | Amp |
| pMT013 | HiFi-reaction between NcoI-cut pETDuet1 and geneblock carrying RecA (*S. aureus*) sequence / Uniprot A5ISG9 | Amp |
| pMT021 | HiFi-reaction between NcoI-cut pETDuet1 and geneblock carrying RecA (*S. venezuelae*) sequence / Uniprot P48295 | Amp |
| pMT025 | Amplification and re-ligation of pMT011 plasmid using T46/T49 primers. | Amp |
| pNPTS138 | ((*2*)) | Kan |
| pXGFPC-1 | ((*3*)) | Spec |
| pNABC865 | *hupA2* gene (along with ~324 bp upstream) from *C. crescentus* was amplified using primers DR16/DR17. A fragment encoding a 20-amino-acid linker fused to mCherry was amplified using primers DR09/DR10. The downstream ~600 bp region of *hupA2* was amplified using primers DR18/DR19. These fragments were assembled into BamHI- and NheI-linearized pNPTS138 using Gibson assembly to generate a C-terminal *hupA2*-mCherry fusion construct. | Kan |
| pNABC1024 | ([4]) | Tet |
| pNABC1026 | ([4]) | Tet |

| | | |
|---|---|---|
| pNABC1030 | The last ~600 bp of CCNA_01879 was amplified using primers AB_239 and AB_248. The tetracycline resistance cassette was amplified using primers AB_184 and AB_185, and the ~600 bp region downstream of CCNA_01879 was amplified using primers AB_242 and AB_243. These fragments were fused by splicing by overlap extension (SOE) PCR using primers AB_239 and AB_243. The resulting fragment was digested with BamHI and NheI and ligated into BamHI–NheI-digested pNPTS138. The construct was verified by sequencing. | Tet |
| pNABC1032 | The full-length *recA* gene from *Caulobacter crescentus* was amplified using primers AC_180 and AC_202, digested with NdeI and KpnI, and ligated into NdeI–KpnI-digested pXGFC-1 to generate a C-terminal GFP fusion to RecA. A 20-amino-acid linker was included between RecA and GFP. | Spec |

**Table S4. Plasmids used in this study.**

| Strain | Genotype | Construction details / Source |
|---|---|---|
| NABC413 | $P_{lacI}$-*lacI (hfa locus)*; $P_{lac}$-*dnaA (dnaA locus)* | ((*5*)) |
| NABC1025 | $P_{lacI}$-*lacI (hfa locus)*; $P_{lac}$-*dnaA (dnaA locus)*; I-SceI site (after CCNA_00029)::$tet^R$ | pNABC1024 was transformed into NABC413 through a two-step recombination protocol. |
| NABC467 | $P_{lacI}$-*lacI (hfa locus)*; $P_{lac}$-*dnaA (dnaA locus)*; I-SceI site (after CCNA_00029)::$tet^R$ ; $P_{van}$-*ISceI*::$chlor^R$ ; *mipZ-mCherry*::$kan^R$ ; $P_{xyl}$-*recA-YFP*::$spec^R$ | ((*6*)) |
| NABC1027 | CB15N; I-SceI site (after CCNA_00201)::$tet^R$ | pNABC1026 was transformed into CB15N through a two-step recombination protocol. |
| NABC1028 | $P_{lacI}$-*lacI (hfa locus)*; $P_{lac}$-*dnaA (dnaA locus)*; I-SceI site (after CCNA_00201)::$tet^R$ | NABC413 was transduced with lysate of NABC1027. |
| NABC1029 | $P_{lacI}$-*lacI (hfa locus)*; $P_{lac}$-*dnaA (dnaA locus)*; I-SceI site (after CCNA_00201)::$tet^R$; $P_{van}$-*ISceI*::$chlor^R$; $P_{xyl}$-*recA-YFP*::$spec^R$ | NABC1028 was transduced with lysate of a strain harbouring *recA-YFP*::$spec^R$ integrated at the $P_{xyl}$ locus. The resulting strain was transduced with lysate of a strain harbouring I-SceI::$chlor^R$ at the $P_{van}$ locus. |
| NABC475 | $P_{lacI}$-*lacI (hfa locus)*; $P_{lac}$-*dnaA (dnaA locus)*; I-SceI site (after CCNA_00727)::tetR ; $P_{van}$-*ISceI*::$chlor^R$ ; $P_{xyl}$-*recA-YFP*::$spec^R$ | ((*6*)) |
| NABC1031 | $P_{lacI}$-*lacI (hfa locus)*; $P_{lac}$-*dnaA (dnaA locus)*; I-SceI site (after CCNA_01879)::$tet^R$; $P_{van}$-*ISceI*::$chlor^R$; $P_{xyl}$-*recA-YFP*::$spec^R$ | I-SceI site with tetR marker was inserted between CCNA_01879 and CCNA_01880 using two-step recombination with pNABC1030 transformed into NABC413. The resulting strain was transduced with lysate of a strain harbouring *recA-* |

| | | |
|---|---|---|
| | | *YFP::spec*$^R$ integrated at the $P_{xyl}$ locus. Finally, this strain was transduced with lysate of a strain harbouring I-SceI::*chlor*$^R$ at the $P_{van}$ locus. |
| NABC845 | $P_{lacI}$-*lacI (hfa locus)*; $P_{lac}$-*dnaA (dnaA locus)*; I-SceI site (after CCNA_00029)::*tet*$^R$; $P_{xyl}$-*recA-GFP*::*spec*$^R$ | pNABC1032 was transformed into NABC1025 through single crossover reaction. |
| NABC1033 | $P_{lacI}$-*lacI (hfa locus)*; $P_{lac}$-*dnaA (dnaA locus)*; I-SceI site (after CCNA_00029)::*tet*$^R$; *hupA2-mCherry*; $P_{xyl}$-*recA-GFP*::*spec*$^R$ | pNABC865 was transformed into NABC845 through two step recombination procedure. |
| NABC1034 | $P_{lacI}$-*lacI (hfa locus)*; $P_{lac}$-*dnaA (dnaA locus)*; I-SceI site (after CCNA_00201)::*tet*$^R$; $P_{van}$-*ISceI*::*chlor*$^R$; *parS*$^{pMT1}$site (after CCNA_00186)::*spec*$^R$; $P_{xyl}$-*YFPparB*$^{pMT1}$*,mCherry-parB*$^{P1}$*::kan*$^R$ | ((*4*)) |
| NABC1035 | $P_{lacI}$-*lacI (hfa locus)*; $P_{lac}$-*dnaA (dnaA locus)*; I-SceI site (after CCNA_00727)::*tet*$^R$; $P_{van}$-*ISceI*::*chlor*$^R$; *parS*$^{pMT1}$site (after CCNA_00747)::*spec*$^R$; $P_{xyl}$-*YFPparB*$^{pMT1}$*,mCherry-parB*$^{P1}$*::kan*$^R$ | ((*4*)) |

**Table S5. *C. crescentus* strains used in this study.**
